# Genome shuffling enables quantitative trait locus mapping in *Bacillus subtilis*

**DOI:** 10.1101/2025.07.14.664384

**Authors:** Delyana P. Vasileva, Hari B. Chhetri, Leah H. Hochanadel, Jared C. Streich, John H. Lagergren, Matthew J. Lane, Edanur Oksuz, Dana L. Carper, Benjamin Rubino, Sadie Chitwood, Zachary D. Schmitz, Dawn M. Klingeman, Paul E. Abraham, Richard J. Giannone, Daniel A. Jacobson, Joshua K. Michener

**Affiliations:** Biosciences Division, Oak Ridge National Laboratory; Oak Ridge, TN, 37830, USA; Center for Bioenergy Innovation, Oak Ridge National Laboratory; Oak Ridge, TN, 37830, USA; Bredesen Center for Interdisciplinary Research and Graduate Education, University of Tennessee, Knoxville; Knoxville, TN, 37996, USA

## Abstract

Genetic mapping is a powerful tool for eukaryotic genetics that has only been applied to bacteria in limited circumstances. Quantitative trait locus (QTL) mapping generally relies on sexual recombination to break linkages between genes, yet bacteria rarely undergo sufficient homologous recombination to generate suitable mapping populations. In this work, we used iterative biparental genome shuffling by protoplast fusion in *Bacillus subtilis* to generate a population of bacteria with substantial random recombination throughout their genomes. Individual shuffled progeny were arrayed in well plates, resequenced, and characterized for a range of complex phenotypes including spore germination and swarming motility. Genetic mapping of the resulting phenotypes identified high-confidence QTLs of moderate size (∼10 kb), and these associations were validated through targeted genetic swaps. This *B. subtilis* QTL population can easily be used to map additional phenotypes, and the general approach for QTL mapping is applicable in a wide range of bacteria.

**One-Sentence Summary:** Bacterial genome shuffling mimics sexual recombination and enables genetic mapping of complex phenotypes to causal alleles.

## Main text

Advances in next-generation DNA sequencing technologies have provided an unprecedented view of the vast genetic landscape of bacteria. However, linking DNA sequences to observable physical traits remains challenging. Even in the best-studied model bacteria, many genes have unknown functions (*1*), and little is known about the genetic networks underlying complex phenotypes or the functional effects of natural sequence variation in bacterial genes and regulatory elements (*2*). Arrayed and pooled genetic screens have revolutionized bacterial functional genomics and identified genetic determinants underlying multiple phenotypes (*3–8)*. However, these approaches are limited to inferring gene function based on loss-of-function or gain-of-function experiments and fail to account for more subtle genetic variation or to capture the complex genetic architecture of quantitative polygenic traits.

Genome-wide association studies (GWAS) and QTL mapping are highly effective techniques in eukaryotic genetics that link genetic variation to phenotypic traits across natural and experimental segregating populations, respectively. Most commonly applied in macroscopic sexually-recombining eukaryotes such as humans and plants, these methods have provided substantial insight into the genetic basis of complex phenotypes and uncovered multiple novel biological mechanisms (*9, 10*). Similar approaches have been developed in yeast to enable fine-scale genetic mapping and analysis of epistatic interactions (*11, 12*). Unfortunately, the use of quantitative genetic mapping techniques in bacteria has been limited by the lack of native systems for sexual recombination and the generally low recombination rates within bacterial populations (*13*). While the GWAS approach has been successfully applied to extant bacterial populations in several instances, these studies have predominantly focused on pathogenic species with large collections of clinical isolates that enabled the assembly of suitably diverse mapping populations (*14–16*). Even then, the mapping approach was most successful when applied to genetic variants that arise under strong selection pressure in recombination hotspots, including variants linked to antibiotic resistance (*15*) and virulence (*17*). Since isolating non-pathogens in sufficient numbers is difficult and most bacteria do not undergo sufficient recombination for GWAS, QTL mapping would be more broadly applicable. Genetic mapping with experimentally generated inter-strain recombinants has been attempted on a small scale in some specific cases, where chromosome regions can be transferred between strains through unconventional conjugal mechanisms (*18*). However, a comprehensive experimental framework for QTL mapping is lacking in bacteria.

We have previously shown that genome shuffling by protoplast fusion between genetically diverse *Bacillus* strains generates frequent, unbiased, genome-wide recombination that mimics the effects of sexual recombination (*19*). In this study, we leveraged protoplast fusion to establish a bacterial QTL mapping platform (Fig. 1A-F). We demonstrated the utility of this approach by identifying causal genetic loci with high statistical power, as well as its potential to reveal the genetic basis of a broad range of complex phenotypes.

**Fig. 1.**
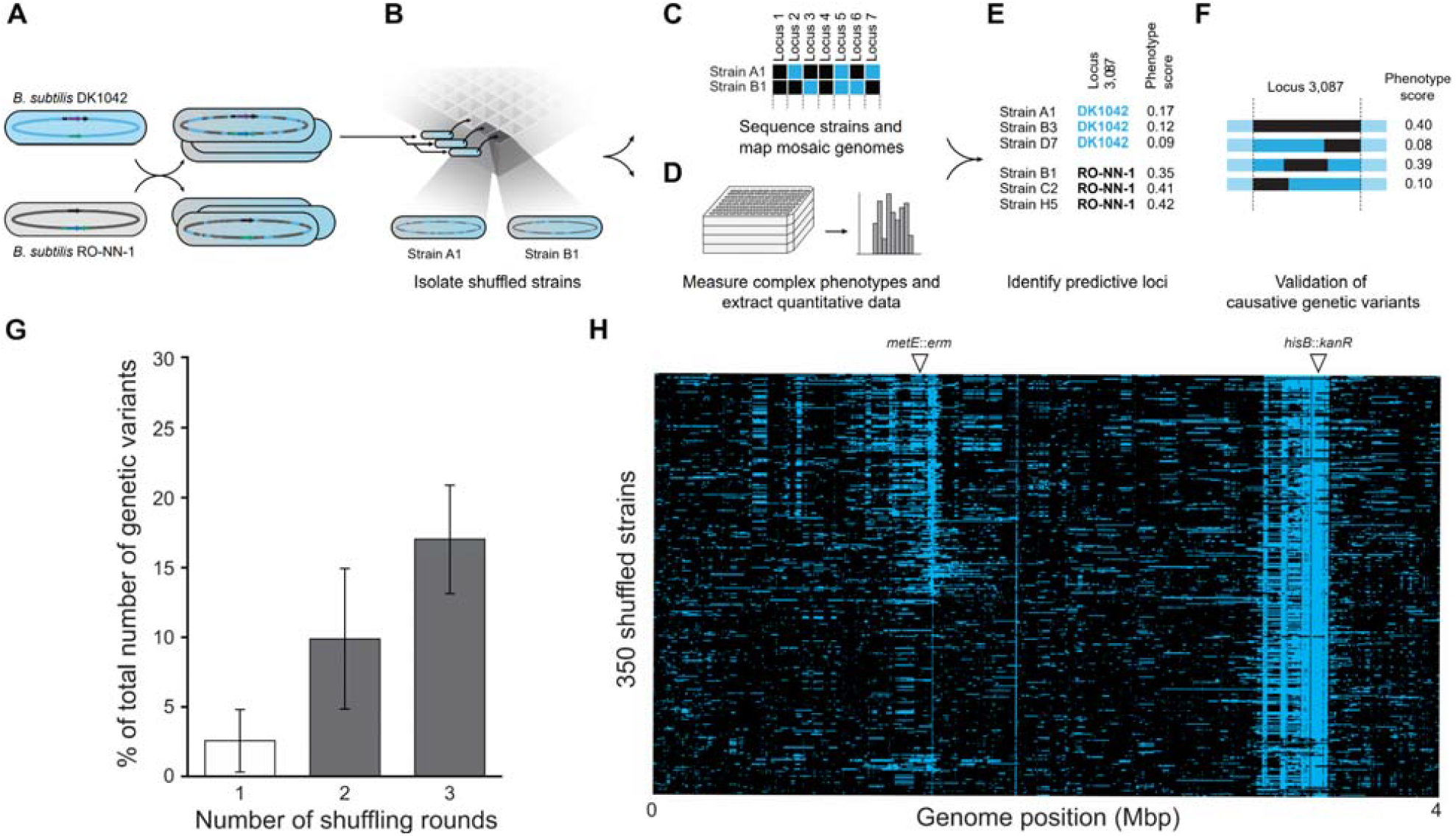
Overview of bacterial QTL mapping. (**A**) Replacing amino acid biosynthesis genes with antibiotic resistance markers in genetically divergent parents allows for selection of recombinant hybrid strains containing different marker allele combinations following genome shuffling by protoplast fusion (fig. S1). (**B**) Repeated rounds of shuffling increase genetic diversity generating mapping populations of highly recombinant progeny. (**C**) The progeny are sequenced to identify genetic contributions from each parent and (**D**) phenotyped. (**E**) Statistical mapping methods are then used to identify genetic loci that correlate with a phenotype of interest. (**F**) These correlations are experimentally validated through CRISPR-mediated markerless genetic swaps. (**G**) The fractions of recombined genome after one round of shuffling between *B. subtilis* RO-NN-1 and 168 (*19*) (white bar) and after two and three rounds of shuffling between strains RO-NN-1 and DK1042 (gray bars) are shown. The number of genetic variants was determined in 24 randomly selected recombinant strains from each shuffle. Error bars represent standard deviation. (**H**) A fine-scale recombination map of the *B. subtilis* QTL population. Each row represents a sequenced individual strain from the 350-strain mapping panel, with blue bars indicating recombined fragments originating from strain DK1042 and black regions representing the genomic portions inherited from strain RO-NN-1. Arrows in panel H indicate locations of the selection markers *metE*::*erm* and *hisB*::*kanR*.

### Development of a highly-recombinant bacterial QTL mapping panel

Our previous work showed that a single round of genome shuffling between pairs of *Bacillus* strains generates recombinant strains with genomes primarily derived from one of the parents but containing multiple unselected DNA fragments originating from the second parent that are randomly recombined throughout the chromosome (*19*). For example, when *B. subtilis* strains 168 and RO-NN-1 were shuffled, DNA fragments from *B. subtilis* 168 replaced, on average, 2.6% of the genome of *B. subtilis* RO-NN-1 (Fig. 1G) and the sizes of the recombined regions were broadly distributed including short inserts (< 100 bp) and large multigene fragments (> 60 kb) (*19*). These two strains vary phenotypically (*20*) and differ in their core genomes at 63,641 single nucleotide polymorphisms (SNPs) (∼98% nucleotide identity), averaging ∼1 SNP every 50 bp. We therefore reasoned that a high frequency of recombination between the chromosomes of strains with this level of genetic divergence would generate in the progeny sufficient admixture of both core genes and accessory pathways to potentially drive mappable variation in phenotypes of interest.

To construct a mapping population, we selected *B. subtilis* RO-NN-1 as the first parent due to its success at genome shuffling in our previous work, and DK1042 as the second parent, a genetically competent strain that exhibits phenotypes lost in the domesticated strain 168 (*21*).

The statistical confidence to identify causal loci is determined by the degree of recombination and the resulting decrease in linkage between genetic variants in the mapping population. To reduce linkage between genetic loci, we aimed to maximize recombination through repeated cycles of genome shuffling using a backcrossing approach (fig. S1). We used three and four rounds of shuffling to increase genome-wide recombination (Fig. 1G and fig. S3) and determined the DNA inheritance patterns of 623 recombinant strains from these shuffles. After removing nearly-identical strains and performing quality control, we retained a total of 350 strains with 60,937 SNPs as a mapping population (Fig. 1H and fig. S4).

While our mapping population used only two parents, multiparental crosses have been applied to improve the power and resolution of QTL studies in eukaryotic genetics by introducing a broader spectrum of genetic and phenotypic variation (*22*). To test whether a similar multiparental approach could be applied in bacterial QTL mapping populations, we shuffled three genetically-divergent *Bacillus* strains, *B. subtilis* RO-NN-1, *B. subtilis* DK1042, and *B. spizizenii* TU-B-10 (fig. S5). The resulting progeny showed diverse inheritance patterns with multiple recombination events from all three parents, demonstrating that a more complex mapping population could be constructed if desired.

### Bacterial QTL mapping identifies causal genetic variants in coding regions and regulatory elements

We next tested whether the mapping population could be used to identify the causal alleles responsible for complex phenotypes. Spore germination is an important developmental phenotype with direct implications for the food industry and human disease (*23*). Previous studies have identified variation in spore germination between *B. subtilis* RO-NN-1 and 168 (*20*), prompting us to use this trait as an initial test case for QTL mapping. To analyze the heritability of spore germination, we generated spores from the two QTL parents and the full panel of shuffled strains and measured their response to L-alanine as the sole germinant. As previously described, we observed significant differences in the spore germination abilities of the two parents (Fig. 2A and fig. S7B). The recombinant strains exhibited a high degree of phenotypic variability with a bimodal distribution (Fig. 2B and fig. S8). We therefore performed QTL mapping using spore germination as a binary phenotype. This analysis identified a ∼10 kb high-confidence locus of strong effect (*H*^*2*^ = 0.94) and the most significant genetic variants were located within a three-gene operon encoding the germinant receptor GerA (Fig. 2C and fig. S9) (*24, 25*). The *gerA* operons of the two parental strains contained 166 sequence variants, among which 46 were non-synonymous (figs. S10 and S11). Analysis of the *gerA* region in the recombinant progeny showed that DK1042 variants were more prevalent in shuffled strains that germinate with L-alanine, whereas RO-NN-1 variants were more common in those that did not germinate, demonstrating the significance of the genotype-trait association (fig. S9).

**Fig. 2.**
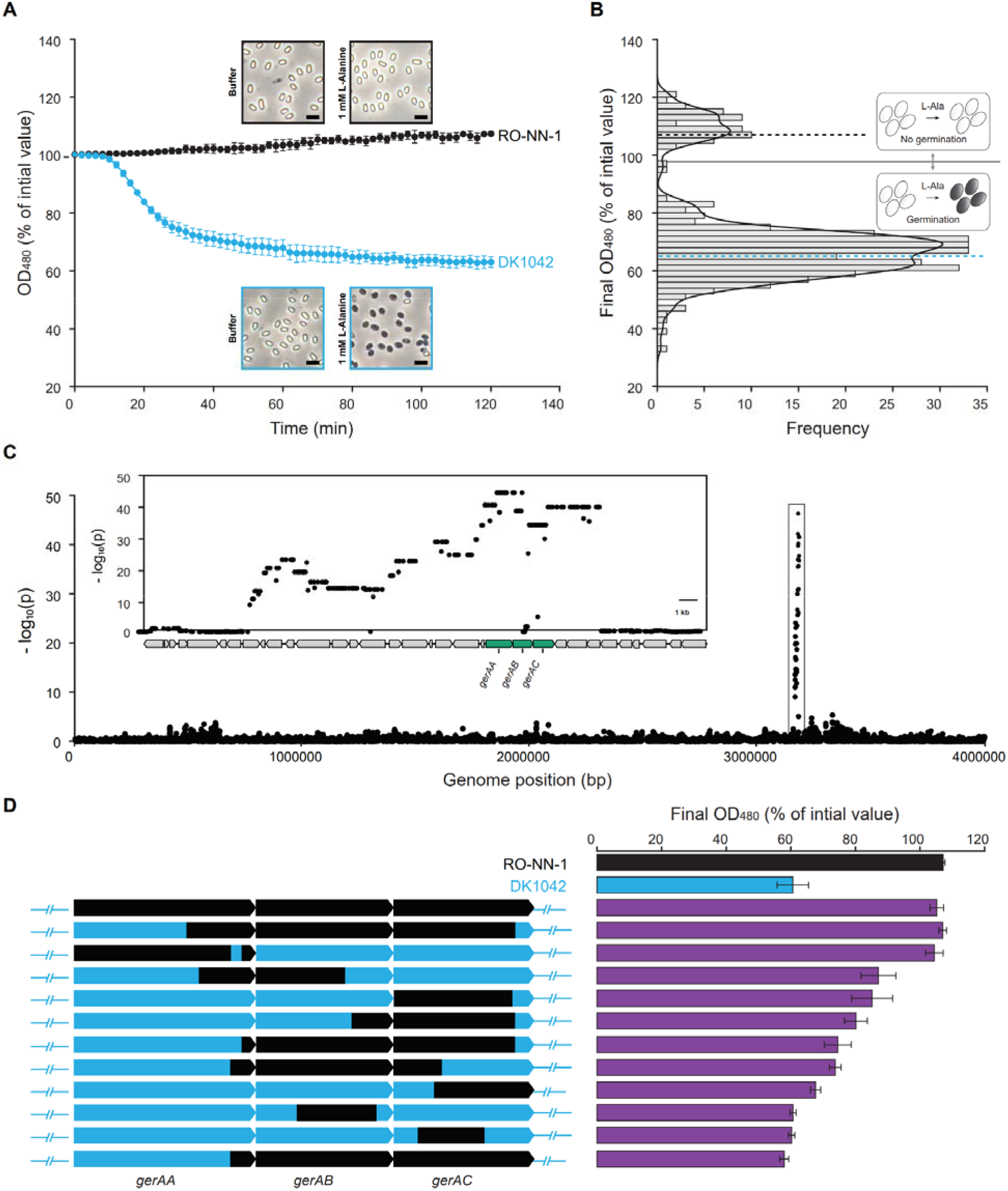
Bacterial QTL mapping identifies genetic variants associated with spore germination. (**A**) *B. subtilis* DK1042 and RO-NN-1 exhibit different spore germination abilities with L-Alanine. Spores of the two strains were incubated with 1 mM L-Alanine and the decrease in optical density was monitored as phase-bright spores transitioned to dark-phase germinated spores. Similar results were obtained with 50 mM L-Alanine (fig. S7B). (**B**) Distribution of spore germination phenotypes across the QTL mapping population. A histogram of optical densities after incubation of spores from each strain with 1 mM L-Alanine for 120 min is shown (fig. S8). Black and blue dashed lines indicate the spore germination data from RO-NN-1 and DK1042, respectively. The solid black line indicates the threshold we defined for spore germination. (**C**) Manhattan plot summarizing the statistical significance of associations between each genetic variant along the genome and spore germination across the *B. subtilis* QTL mapping population. The inset shows a zoomed-in view of the region containing the spore germination QTL. Candidate causal genetic loci from the *gerA* operon are highlighted in green. (**D**) Genetic swaps validate causal genetic loci. Gene segments within the *gerA* operon in strain DK1042 were replaced with corresponding segments from strain RO-NN-1 using a CRISPR-based approach and their spore germination abilities were evaluated. The bar plot shows optical densities measured after incubation of spores of each strain with 1 mM L-Alanine for 120 min. Data from the full library of gene-swapped strains is shown in fig. S12. Black regions indicate genetic fragments originating from strain RO-NN-1 and blue regions indicate segments from strain DK1042. The data represents the mean results of three biological replicates. Error bars represent standard deviation.

To validate our QTL inference, we performed allele replacement experiments by swapping regions within the DK1042 *gerA* operon with corresponding genetic fragments from strain RO-NN-1 and evaluated spore germination of the resulting strains in response to L-alanine (Fig. 2D, and figs. S12 and S13). Spores derived from a strain where the entire DK1042 *gerA* operon was replaced with the corresponding RO-NN-1 allele failed to germinate under these conditions, in a manner similar to RO-NN-1. Strains harboring different *gerA* regions derived from RO-NN-1 showed diverse quantitative phenotypes, including germination rates similar to the QTL parents as well as intermediate phenotypes. Specifically, genetic variants located in the genes encoding the GerAA and GerAC subunits of the germination receptor, as well as interactions between these variants, were associated with different spore germination rates (fig. S12). In contrast to the response with L-alanine, spores of strain RO-NN-1 germinated in response to L-valine in a GerA-dependent manner (figs. S14 and S15), indicating that both strains produce active GerA but with differing sensitivity to common germinants.

By identifying SNPs within functional genes that affect complex phenotypes, we demonstrated that QTL mapping provides a framework to characterize the complex biological effects of natural genetic variation in bacteria beyond simple loss-of-function mutations. Targeted genetic swaps can be applied to narrow down the range of candidate genetic variants in a QTL region and distinguish the effects of SNPs within causal genes. Although the majority of the shuffled strains in our QTL population contained either DK1042 or RO-NN-1 variants in the *gerA* locus, seven strains harbored single variants originating from strain DK1042 or different combinations of variants from the two parents. With a larger sample size, the density of recombination events achieved with our cross design has the potential to facilitate fine-scale mapping without the need for additional experimentation to resolve composite QTLs.

We next tested the performance of our QTL platform using an additional phenotype, swarming motility -a model multicellular trait that varies substantially among different *Bacillus* isolates (fig. S16) (*26*). Our analysis revealed significant differences in the swarming motility of the two QTL parents (Fig. 3A and B), and the QTL population displayed a broad continuous distribution of this phenotype (Fig. 3C). Consistent with the spore germination test case, we mapped this trait and identified a highly significant QTL (*H*^*2*^ = 0.66) of moderate size (∼10 kb) associated with swarming motility (Fig. 3D and fig. S19). We then systematically screened loci within the QTL using our allele replacement approach (Fig. 3E and fig. S20). These experiments narrowed the candidate causal variants down to two SNPs: one located in the regulatory region of the transcriptional regulator *rghR*, and the other a synonymous mutation within the coding sequence of *rghR*. A modified DK1042 strain containing the two candidate causal SNPs showed reduced swarming motility, similar to that of the RO-NN-1 strain (Fig. 3E). RghR is known to repress the expression of *rapG* and *rapH*, which encode inhibitors of the surfactin biosynthetic cluster *srfA* (*27*). Both wild-type RO-NN-1 and the DK1042 strain containing the two candidate causal SNPs exhibited significantly lower expression of the RghR and SrfA proteins, as well as reduced surfactin production, compared to strain DK1042 (fig. S21, A and B). Furthermore, addition of surfactin to the growth medium enabled swarming motility in strain RO-NN-1 (fig. S21, C and D), suggesting that variation in surfactin levels is responsible for the phenotypic differences.

**Fig. 3.**
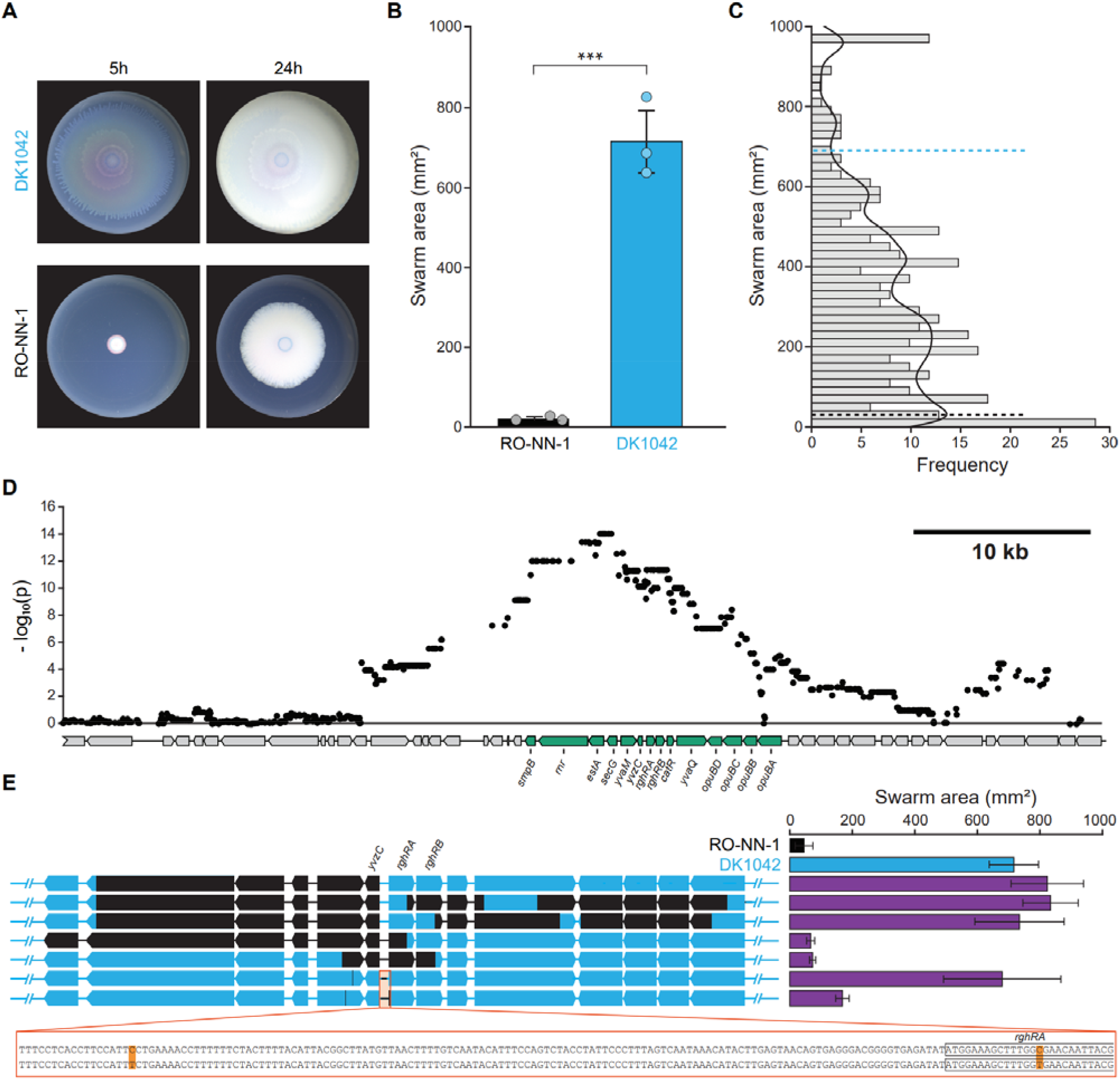
Regulatory genetic variants drive phenotypic differences in swarming motility. (**A**) Comparison of swarming motility of *B. subtilis* DK1042 and RO-NN-1 over the course of 5h and 24h. Representative data from three biological experiments is shown. (**B**) Areas covered by the DK1042 and RO-NN-1 swarms measured after 5h of incubation. The data were analyzed by means of two-tailed Student’s *t* test and represents the average results of three biological replicates. Error bars indicate standard deviation. *p* value < 0.001 (***) is indicated. (**C**) Distribution of swarming motility phenotypes across the QTL population. A histogram of swarm areas after incubation for 5h is shown. (**D**) Inset of a Manhattan plot (fig. S19) shows a 50-kb genomic region including the QTL for swarming motility. Candidate causal genetic loci are highlighted in green. (**E**) Genetic swaps within the QTL validate the association and identify two candidate causal genetic variants. Black regions indicate genetic fragments originating from strain RO-NN-1 and blue regions indicate segments from strain DK1042.

These results highlight that our QTL mapping approach can uncover subtle regulatory mechanisms and genetic factors contributing to phenotypes related to public goods production, which would likely be overlooked in traditional pooled genetic screens.

### Genome shuffling creates mappable variation across a wide range of bacterial traits

To explore further the applicability of our platform for mapping a variety of traits, we selected the 23 most genetically diverse shuffled strains from the QTL population and evaluated phenotypic variability across this panel (Fig. 4). We tested both phenotypes relevant to microbial ecology and traits of interest for bacterial engineering. Since none of the shuffled strains in the QTL population carried the DK1042-derived plasmid pBS32 and the effects of this plasmid on some of the tested phenotypes is not known, we used a cured strain as the parental control in these experiments.

**Fig. 4.**
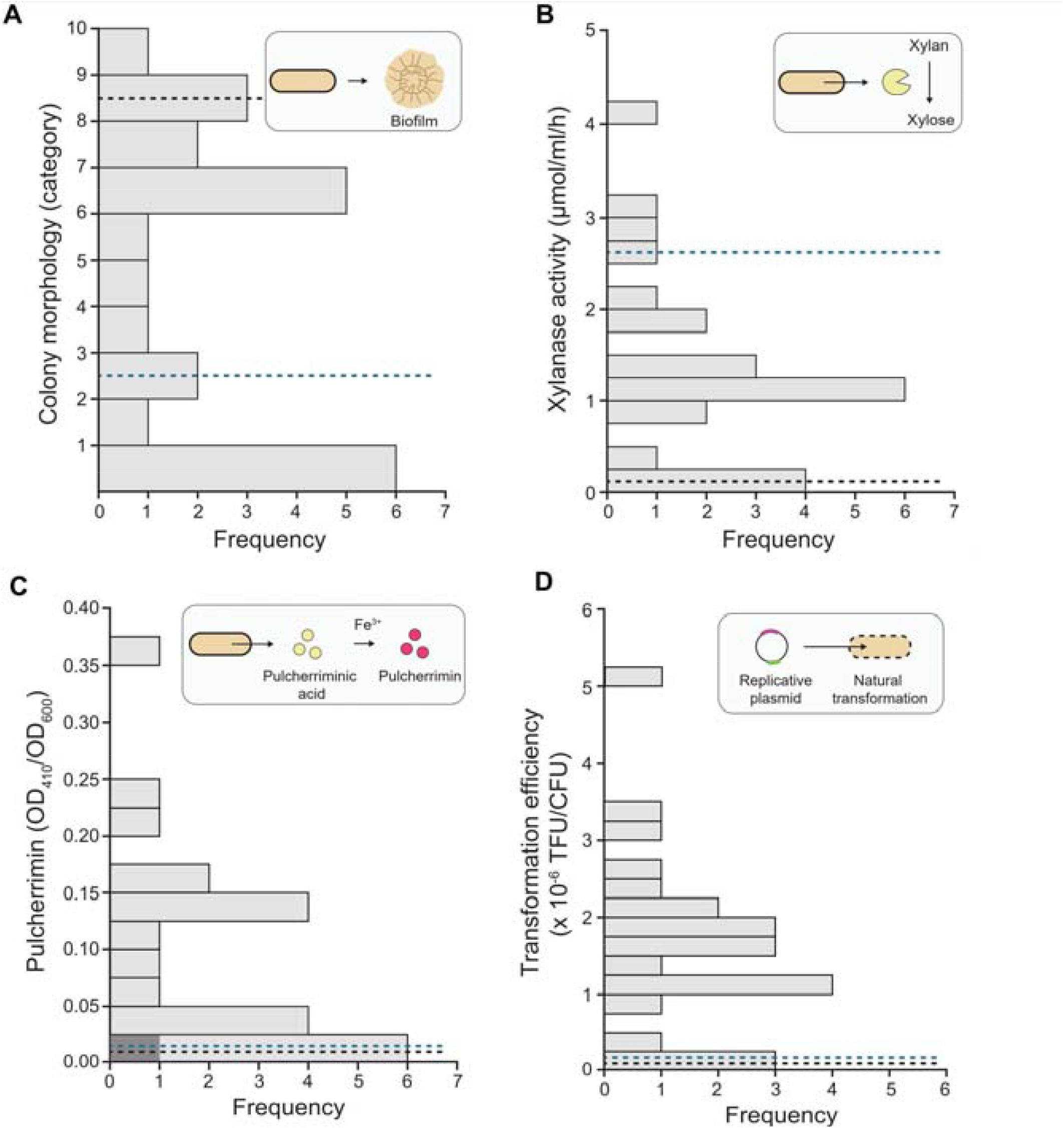
Bacterial QTL mapping is applicable to a wide range of complex phenotypes. Phenotypic variability was tested across the 23 most genetically diverse strains from the QTL population for a variety of traits. (**A**) Distribution of colony morphology phenotypes under biofilm-promoting conditions. A histogram summarizing colony morphology category frequencies (fig. S22) is shown. (**B**) Distribution of extracellular xylanase activity phenotypes. A histogram of xylanase activities (fig. S23) is shown. (**C**) Distribution of pulcherrimin biosynthesis phenotypes. A histogram of pulcherrimin production (fig. S24) is shown. All strains except RO-NN-1 and one shuffled strain indicated by a dark gray bar carried the pulcherrimin biosynthesis cluster. (**D**) Distribution of transformation efficiency phenotypes. A histogram of transformation efficiencies via natural competency (fig. S25) is shown. Dark blue dashed lines indicate phenotypic data from RO-NN-1 and the cured DK1042 strain JM300 (Materials and Methods), respectively.

Biofilm formation is a complex phenotype that involves multiple genes and genetic interactions (*28*). We tested biofilm architectures across the 23-strain panel and the parental controls and categorized the colonies based on their morphology (Fig. 4A and fig. S22). Our analysis revealed that genome shuffling generated extensive variability in colony patterns. In addition, we evaluated enzyme activity and secondary metabolite production phenotypes, including extracellular xylanase activity (Fig. 4B and fig. S23A) and biosynthesis of pulcherrimin, an iron-binding reddish pigment (Fig. 4C and fig. S24). Finally, we assessed phenotypes relevant to genetic engineering, including promoter leakiness (fig. S23B) and transformation efficiency (Fig. 4D and fig. S24). All of these assays also revealed substantial phenotypic variability among the 23 shuffled strains. Across the different assays, we frequently identified shuffled strains exhibiting phenotypes that are not intermediate of the parental strains, but rather represent quantitatively or qualitatively new phenotypes. Collectively, these findings indicated that the tested traits are complex and that the progeny represent a new mixture of alleles, enabling the discovery and mapping of novel phenotypes.

## Conclusions

The power of phenotype-genotype association studies in bacteria is limited by the relatively low recombination rates in natural bacterial populations. To overcome these limitations, we developed an approach to generate highly recombinant panels, enabling high-confidence genetic mapping. We detected QTLs spanning on average ∼10 kb, though our data suggest that genetic mapping resolution in bacteria can be improved either by increasing the number of recombinant strains in the QTL population or the overall recombination frequency across the genome. Our findings demonstrate that, beyond gene presence or absence, natural genetic variation in both coding and non-coding regions significantly influences bacterial phenotypes. Bacterial QTL mapping, therefore, provides a more complete picture of the genetic architecture underlying complex bacterial traits by systematically exploring the functional impacts of the standing genetic diversity within natural microbial populations.

The iterative protoplast fusion recombination method developed in this study can be extended to a wide range of Gram-positive bacteria. However, its use in Gram-negative bacteria can be potentially challenging due to the complexity of their cell walls, which makes protoplasting more difficult. Large-scale interstrain genetic exchange in Gram-negative bacteria can be achieved through alternative approaches like chromosomal conjugation and natural transformation (*29*– *32)*. These approaches can be leveraged in future efforts to create libraries with sufficient genome-wide recombination for genetic mapping.

We envision that bacterial QTL mapping, in combination with the traditional deletion and pooled genetic screens, would enable a powerful platform technology for comprehensive functional bacterial genomics.

## Acknowledgments

We thank Dr. Ilenne Del Valle for providing us with the plasmid pIDV40 used for the transformation efficiency assays.

## Author contributions

Investigation: DPV, LHH, JCS, EO, DLC, BR, SC, ZDS, DMK Methodology: DPV, HPC, JCS, JHL, MJL, BR, PEA

Software: HPC, JCS, JHL, MJL, PEA

Visualization: DPV, HPC, JCS, JHL, MJL

Conceptualization: DAJ, JKM

Funding acquisition: DAJ, JKM

Project administration: DAJ, JKM Supervision: DAJ, JKM, PEA, RJG

Writing – original draft: DPV, LHH, HBC, JCS, JHL, MJL, EO, DLC, DAJ, JKM Writing – review & editing: SC, ZDS, DMK, PEA, RJG,

