## Supplemental Material for "Genome shuffling enables quantitative trait locus mapping in *Bacillus subtilis*"

**The PDF file includes:**

Materials and Methods
Supplementary Text
Figs. S1 to S26
Tables S1 to S3
References

**Other Supplementary Materials for this manuscript include the following:**

Data S1

### Materials and Methods

#### Strains and culture conditions

Bacterial strains used in this study are listed in Table S1. *Bacillus subtilis* subsp. *subtilis* RO-NN-1 (*1*, *2*), *Bacillus subtilis* subsp. *subtilis* NCIB 3610 (*3*, *4*), *B. subtilis* DK1042 (*5*), BKK34900 (*6*), BKE13180 (*6*), *B. spizizenii* TU-B-10 (*1*, *7*, *8*), *B. velezensis* FZB42 (*9*, *10*), and *B. velezensis* GB03 (*11*) were obtained from the Bacillus Genetic Stock Center (BGSC). *B. subtilis* DSM 1970 was obtained from the German Collection of Microorganisms and Cell Cultures (DSMZ). *Escherichia coli* DH10B and JM109 were used for plasmid construction and propagation. Unless otherwise indicated, *E. coli* and *B. subtilis* strains were grown in Lysogeny broth (LB) (casein peptone 10 g/l, NaCl 10 g/l, yeast extract 5 g/l) or LB broth solidified with 1.5% (w/v) Bacto agar. *E. coli* cultures were grown at 37 ºC and *B. subtilis* cultures were grown at 30 ºC or 37 ºC. Growth media were supplemented with antibiotics, when appropriate, as follows: kanamycin (50 µg/ml), erythromycin (20 µg/ml) plus lincomycin (12.5 µg/ml), tetracycline (15 µg/ml), chloramphenicol (25 µg/ml for *E. coli* and 15 µg/ml for *B. subtilis*), carbenicillin (100 µg/ml), and spectinomycin (100 µg/ml). All oligonucleotides and plasmids used for strain construction and verification are summarized in Tables S2 and S3. Strain construction is described in Supplemental Methods.

#### Construction of a *B. subtilis* biparental mapping population

Genetically diverse recombinant strains for QTL mapping were created by three and four rounds of genome shuffling through protoplast fusion using as parental strains *B. subtilis* RO-NN-1 and DK1042. The process of construction of the QTL population is illustrated in Fig. S1. Protoplasts were prepared, fused, and regenerated as described previously (*12*).

In the first round of shuffling (S1), strain RO-NN-1 was crossed with DK1042 Δ*hisB*::*kan* Δ*metE*::*erm* ('DK1042 HK ME') and strain DK1042 was crossed with RO-NN-1 Δ*hisB*::*kan* Δ*metE*::*erm* ('RO-NN-1 HK ME'). Δ*hisB*::*kan* $metE^+$ ('S1 HK') and Δ*metE*::*erm* $hisB^+$ ('S1 ME') recombinant progeny were selected on minimal medium (MM) agar (*13*) supplemented with histidine (300 µM) and kanamycin and MM agar supplemented with methionine (1 mM), erythromycin and lincomycin, respectively. S1 HK and S1 ME pools were made by scraping all colonies off selective plates. The cells were resuspended in 500 µl ddH$_2$O, 5 µl of each cell suspensions were then inoculated into 5 ml selective MM and grown overnight at 37 ºC. Pools were stored as glycerol stocks. Select individual strains from the S1 HK and S1 ME pools were resequenced and RO-NN-1 was identified as the major contributor to the offspring genomes.

In the second round of shuffling (S2), the S1 HK pool was crossed with DK1042 Δ*metE*::*erm* ('DK1042 ME'), the S1 ME pool was crossed with DK1042 Δ*hisB*::*kan* ('DK1042 HK'), and the S1 HK was crossed with S1 ME. Prototrophic ($hisB^+$ $metE^+$, 'WT') and double-resistant progeny (Δ*hisB*::*kan* Δ*metE*::*erm*, 'HK ME') from each cross were selected on MM agar and LB agar supplemented with kanamycin and erythromycin, respectively. WT and HK ME pools were created from each cross as described above and stored as glycerol stocks. Individual strains from each pool were resequenced. The number of genetic variants per strain was estimated using the Single nucleotide polymorphism (SNP) call function of Geneious Prime (versions 2019.0 and 2024.0) and the RO-NN-1 reference genome. S2 WT and S2 HK ME strains resulting from the S1 ME x DK1042 HK cross showed the most genetic diversity and were selected for the next round of shuffling.

In the third round of shuffling (S3), the S2 WT pool was crossed with DK1042 HK ME giving S3 HK1 and S3 ME1 recombinant progeny and the S2 HK ME pool was crossed with DK1042 giving S3 HK2 and S3 ME2 shuffled strains. All S3 HK1 and S3 HK2 strains were combined to create the S3 HK pool.

In the fourth round of shuffling (S4), the S3 HK pool was crossed with strain RO-NN-1 ME giving S4 HK ME recombinant progeny.

Five hundred seventy-three individual S3 HK1, S3 HK2 and S3 ME1 strains and 50 S4 HK ME strains were arrayed in 96-deep-well plates and their genomes were resequenced. SNPs were identified and boundaries of the genomic fragments inherited from each parent were determined as described below. Nearly identical strains, rare genetic variants and plate specific variants were filtered out resulting in a total of 350 genetically diverse strains. Detailed analyses of the genome sequences of these strains revealed that approximately 10% of the wells did not contain pure cultures. Cultures from all 350 wells were streaked to single colonies on selective plates. Individual strains were arrayed in 96-deep-well plates (fig. S2) and their genomes were resequenced.

**Triparental genome shuffling**

Triparental crosses were carried out by protoplast fusion as described above. A pool of 24 S4 HK ME strains from the RO-NN-1 x DK1042 biparental cross was shuffled with wild-type *B. spizizenii* TU-B-10. Strains from both potential recombinant genotypes, either Δ*hisB*::*kan metE*$^{+}$ (‘HK’) or Δ*met*::*erm hisB*$^{+}$ (‘ME’), were selected on MM agar supplemented with histidine (300 μM) and kanamycin and MM agar supplemented with methionine (1 mM), erythromycin and lincomycin, respectively. Individual recombinant strains were isolated and resequenced.

**Sequencing**

Genomic DNA from the *B. subtilis* QTL panel was extracted using the Mag-Bind® Bacterial DNA 96 Kit (Omega Bio-Tek, Norcross, GA, USA) following the manufacturer’s instructions. DNA was then sequenced by the Joint Genome Institute (JGI). The whole-genome sequencing data have been submitted to NCBI Sequence Read Archive and assigned the identifier ID.

**Variant calling**

Variant calling for the whole-genome resequencing data was performed as described previously (*12*). Briefly, raw sequence data with an average sequencing depth of 80x to 100x were processed by trimming low-quality bases based on Phred quality scores using Trimmomatic such that any leading or trailing sequence bases of the sequence read with Phred quality score less than 3 were removed and the read less than 70 bases long were discarded (Trimmomatic version 0.39; (*14*)). The trimmed reads were aligned to the reference genome using the BWA-MEM algorithm (BWA version 0.7.17; (*15*)), and the resulting alignments were converted to compressed BAM files using SAMtools (SAMtools version 1.3.1; (*16*)). Genomic coordinates were sorted using Picard Tools (Picard version 2.27.4; Broad Institute, 2019) and indexed with SAMtools (*16*). SNPs were called using the GATK HaplotypeCaller (GATK version 4.1.8.1; (*17*)). Hard-filtering criteria, as recommended by the GATK developers, were applied to remove low-quality and potentially biased variants. The following filters were used: QD < 2.0, QUAL < 30.0, SOR > 3.0, FS > 60.0, MQ < 40.0, MQRankSum < -12.5, and ReadPosRankSum < -8.0. Variants failing to meet these criteria were excluded from further analysis.

Genomes of (RO-NN-1 x DK1042) S4 HK ME x TU-B-10 strains were analyzed using Geneious Prime (version 2024.0.7). Sequence reads were mapped to the RO-NN-1, NCIB 3610, and TU-B-10 reference genomes. SNPs were identified and boundaries of the genomic fragments inherited from each parent were determined.

**Phenotypic Data Preparation and Heritability Estimation**

Phenotypic data were initially screened for outliers using the Median Absolute Deviation (MAD) method. Any data points with a MAD distance greater than 6 were classified as outliers and excluded from all subsequent analyses.

To assess the genetic control of traits and the repeatability of measurements, broad-sense heritabilities ($H^2$) were estimated. The heritability was calculated using the following formula:

$$H^2 = \frac{\sigma_G^2}{\sigma_G^2 + \sigma_\epsilon^2}$$

where $\sigma_G^2$ represents the genotypic variance, attributed to differences between bacterial strains, and $\sigma_\epsilon^2$ represents the residual variance.

Additionally, to adjust for potential confounding due to temporal variation, the time at which measurements were conducted was included as a covariate in the statistical model to account for batch effects.

The random effects of bacterial strains (genotypes) were modeled using Best Linear Unbiased Predictors (BLUPs) for each phenotypic trait. These BLUPs were subsequently used in all genome-wide QTL analyses.

**Heritability Visualization**

Genetic maps of shuffled strains were derived using a python codebase that can be found at https://github.com/LaneMatthewJ/heritability_illustrator/. In the python codebase, the code was designed to analyze a user-specified region of genetic data filtered by a start and stop position within the genome.

The supplied VCF file was read in as a pandas dataframe (*18*, *19*). All SNPs between the loci range of 0 and 4.2M were selected, such that all SNPs within the genome were maintained. All genomic information was transformed to a "dominant match" boolean, such that if the SNP in the progeny sample matched the SNP in the dominant parent, the value was converted to 1, otherwise 0. No samples were clustered. All unique samples were plotted as a heatmap using the Seaborn heatmap plot such that each row denotes individual samples, and each column a SNP (*20*).

**QTL mapping**

Association mapping on the phenotypic traits was conducted using the following mixed linear model (MLM) implemented in the GAPIT package in R (*21*):

$$Y = X\beta + Zu + \epsilon$$

where, $Y$ is the $n \times 1$ vector of phenotypic values, with $n$ sample size, $\beta$ is the $p \times 1$ vector of fixed additive SNP effects, with $X$ as its corresponding n x p incidence matrix of genotypic data, $u$

represents the *n x 1* vector of random effects due to genotypes, which includes the genetic relationship matrix, estimated using Zhang method (*22*), Z is the incidence matrix for random effects, and $\epsilon$ is the *n x 1* vector of residuals accounting for unexplained phenotypic variance.

A total of 60,937 high-quality SNPs with a minor allele frequency of at least 0.03 were used in the association analysis, derived from 350 unrelated bacterial strains. The SNP dataset was filtered to remove highly clonal individuals.

**Spore preparation**
Spores for spore germination assays were routinely produced using the nutrient exhaustion method in liquid Difco sporulation medium (DSM) (*23*). Strains were grown overnight in 96-deep-well plates at 37 ºC in LB broth supplemented with antibiotics where necessary. Cells were then washed with 1X phosphate-buffered saline (PBS, pH 7.4), pellets were resuspended in DSM and cultures were incubated at 37 ºC for 72h. Spores were harvested by centrifugation at 3,000 rpm for 15 min at room temperature and washed five times with 1X PBS buffer. Spore suspensions were stored in 1X PBS at 4 ºC until analysis.

**Spore germination assay**
Spore germination assays were carried out as described previously (*24*), with some modifications. Spores were normalized to an optical density at 480 nm ($OD_{480}$) of 0.6 in $ddH_2O$ and 200 µl spore suspensions were then transferred to clear, flat-bottom, 96-well plates. After heat activation at 70 ºC for 30 min, spores were cooled at 4 ºC for 15 min and harvested by centrifugation at 3,000 rpm for 15 min at room temperature. The supernatants were discarded, and the pellets were resuspended in 10 mM Tris-HCl, 20 mM KCl, pH 7.4 (buffer) or buffer supplemented with nutrient germinants (1 or 50 mM L-alanine, 1 or 50 mM L-valine). Germination assays were carried out at 37 ºC, and the $OD_{480}$ was monitored every 2 min for 2h using a Synergy H1 plate reader (BioTek, Winooski, VT, USA) with Gen 5 software (version 3.03).

**Microscopy**
Spores were concentrated by centrifugation and then immobilized on agarose pads (1.5% agarose in 1X PBS buffer). Phase-contrast microscopy was performed using an Axio Imager.A2 Zeiss microscope (Carl Zeiss, Oberkochen, Germany) equipped with Zeiss Plan-Apochromat 100x/1.4 Oil Ph3 immersion objective lens (Carl Zeiss, Oberkochen, Germany) and Canon EOS Rebel T1i camera (Canon, Tokyo, Japan). Image processing was performed using EOS Utility 2 software (Canon, Tokyo, Japan). For each strain, at least three independent spore preparations were subjected to microscopy and representative images are shown.

**Swarm expansion assay**
Swarming motility assays were conducted as described previously (*25*), with some modifications. Cultures of *B. subtilis* strains were inoculated in LB broth supplemented with antibiotics as needed and grown overnight in a shaking incubator at 37 ºC and 250 rpm. The overnight cultures were subcultured into fresh LB in culture tubes or 96-deep-well plates and grown to mid-log phase ($OD_{600}$ ~0.5) at 37 ºC and 250 rpm. Cells were then harvested by centrifugation and resuspended to an $OD_{600}$ of 0.6 in 1X PBS buffer (pH 7.0) containing 0.5% India ink (Higgins). LB containing 0.7% agar was prepared freshly on the day of the experiment in clear, flat-bottom, 6-well plates (5 ml agar per well) and the plates were dried open-faced in a laminar flow hood for 30 min. One microliter of resuspended cells was spotted onto the center of an agar well, allowed to dry for 20

min and incubated at 37 ºC for 5h. For experiments with exogenous surfactin, surfactin (cat#S3523, Sigma-Aldrich, St. Louis, MO, USA) was dissolved in pure ethanol and added to the swarm agar wells at a final concentration of 12.5 μM.

**Biofilm formation**

Colony biofilm formation of *B. subtilis* strains was induced using MSgg agar medium (5 mM potassium phosphate (pH 7.0), 100 mM MOPS (pH 7.0), 2 mM $MgCl_2$, 700 μM $CaCl_2$, 50 μM $MnCl_2$, 50 μM $FeCl_3$, 1 μM $ZnCl_2$, 2 μM thiamine, 0.5% glycerol, 0.5% glutamate, 1.5% (w/v) Bacto agar) (*26*), supplemented with amino acids for auxotrophic strains. Cultures of *B. subtilis* strains were inoculated in LB broth and grown overnight in a shaking incubator at 37 ºC and 250 rpm. MSgg agar was prepared in clear, flat-bottom, 6-well plates (6 ml agar per well). One microliter of overnight culture was spotted onto the center of an agar well, allowed to dry for 20 min and incubated at 30 ºC for 72h.

**Swarm and biofilm image acquisition**

A custom XY-robot was designed and 3D printed to automate the collection of high-resolution image data of swarm expansion assays and biofilms on 6-well plates. The system used microcontrollers to move clear 6-well laboratory plates under a fixed camera in uniform lighting conditions. Each well was systematically positioned under the camera and illuminated using a dark field approach, in which a strip of white LED lights illuminated the bacterial colony from an angle and did not shine directly into the camera. The system was operated by a computer (connected via USB) and automatically tracked samples using manual-input sample IDs, barcode IDs, and/or radio frequency IDs. The system supports two cameras: first a low resolution camera (Hotpet 8MP Optical 10X Zoom 5-50mm Lens Industrial Web Camera, Hotpet, China) which was used to image the population of *B. subtilis* strains, and was later upgraded to a high resolution camera (Canon EOS 4000D, Canon, Tokyo, Japan) used to conduct the validation of the swarming phenotype.

**Swarm image analysis**

Swarm assay phenotyping was formulated as a semantic segmentation problem, in which manually-annotated *B. subtilis* swarm images were used to train a region growing convolutional neural network (CNN) model based on (*27*) to identify pixels along the boundaries of the swarm (fig. S17). Each image was stored in PNG format with a spatial resolution of 896x1152 pixels. A training set of 60 images was manually annotated using the GNU Image Manipulation Program (*28*) in which we manually traced the boundary of each colony and stored the contour as a black-and-white PNG image. The annotated images and masks were then randomly partitioned into 80% training and 20% validation sets. Each image was further broken into smaller training tiles of size 128x128 which were augmented during training using random flips, rotations, Gaussian blur, and color jitter corresponding to brightness, contrast, hue, and saturation. The CNN model was trained to predict the class labels (i.e., boundary vs background) of the center 3x3 square of each tile using a batch size of 512, the focal loss objective function (*29*), and Adam optimizer (*30*) with default hyperparameters. Early stopping of 20 epochs was used to ensure model convergence and prevent overfitting to the training set. The model weights from the epoch that minimized validation error were selected as the final weights used for inference.

Inference was formulated as a region growing task following (*27*), where CNNs iteratively add pixels to a growing colony boundary segmentation. Swarms with high-contrast boundaries were accurately detected, but swarms with low-contrast and diffuse boundaries (especially close to the edge of the well) could result in false positive and false negative predictions. Thus, we developed a segmentation post-processing procedure that detected the longest continuous boundary from a swarm, and used the segment to compute the best fit circle. Note that for swarms with contiguous boundaries, the whole boundary was used as the line segment. Each circle (parameterized by the radius and the row and column location of the center) was optimized by minimizing the Chamfer distance between the circle and the line segment. The best fit circle was then used to predict the swarming phenotypes (i.e., colony radius and area). A conversion factor based on the size of the 6-well plate was used to convert between pixel units and millimeters. Each result (predicted boundary and best fit circle) was manually inspected for accuracy. Any images with errors were manually corrected and phenotyped.

**Reconstruction of causal genetic variants**

To identify and validate causal genetic variants, we functionally profiled regions within each QTL through CRISPR-Cas9 mediated genetic swaps using pJOE9658.1, which carries the *cas9* gene under the control of tetracycline-inducible *tetLM* riboswitch and a single-guide RNA (sgRNA) encoding sequence transcribed from the strong semisynthetic promoter $P_{VanP*}$ (*31*). Alleles in strain DK1042 were replaced with the alternative variants of these alleles from strain RO-NN-1. 20-nucleotide sgRNA spacer sequences targeting the regions of interest were designed using the in-build feature “Find CRISPR sites” in Geneious Prime (versions 2023.0 and 2024.0). The reference genome of strain RO-NN-1 was used as an off-target database. sgRNA spacer sequences were cloned into the BsaI sites of pJOE9658.1 by ligating annealed oligonucleotides as described previously (*31*, *32*). Oligonucleotides were designed to include sequences that are complementary to the sticky ends generated by BsaI digestion of the plasmid. Repair templates were generated by PCR amplification of suitable regions from RO-NN-1 genomic DNA. The repair templates were then cloned between the SfiI sites of pJOE9658.1 as described previously (*31*, *32*) or cotransformed with the plasmids as PCR products. All pJOE9658.1 derivative constructs were verified by next-generation whole-plasmid sequencing (Plasmidsaurus Inc., Eugene, OR, USA).

Strain DK1042 was transformed with the pJOE9658.1 derivative plasmids and repair templates by natural competency, following standard protocols (*6*). Transformants were plated on LB agar plates supplemented with 50 µg/ml kanamycin and 200 ng/ml tetracycline (to induce Cas9 expression) and incubated for 24h at 30 ºC. The resulting colonies were streaked on LB agar without antibiotics and incubated for 24h at 45 ºC to cure the plasmid. Single colonies were then tested for the loss of the plasmid by patching on LB agar plates with kanamycin. The colonies that failed to grow on these plates were analyzed for the presence of the desired edits using PCR amplification of the targeted region followed by long-read sequencing (Plasmidsaurus Inc., Eugene, OR, USA). Select strains were also verified by whole-genome sequencing on an Illumina MiSeq instrument (Illumina, San Diego, CA) as described previously (*12*).

**Transformation frequency assays**

Competent cells for transformation frequency assays were prepared as described previously (*33*), with some modifications. *B. subtilis* strains were grown overnight in 1 ml LB broth in a shaking incubator at 37 ºC and 250 rpm. Cells were then harvested by centrifugation, washed in 1 ml 1X PBS, and 200 µl cell suspension was inoculated in 3 ml SM1 medium. Following incubation for 3h or for the indicated time period at 37 ºC, 3 ml of SM2 medium was added, and cells were further incubated for 90 min under the same conditions. The cultures were then combined with plasmid DNA at the indicated concentrations and incubated for an additional 1h. Next, 3 ml of LB was added to the cultures and cells were incubated for 30 min. The cells were harvested by centrifugation and resuspended in 300 µl 1X PBS. Serial dilutions were prepared in 1X PBS and cells were plated onto selective LB plates to determine the number of transformants and onto LB plates to determine the number of viable cells. The transformation frequency was determined by calculating the ratio of transforming units to colony-forming units.

**Xylanase activity assay**

Xylanase activity assays were performed as described previously (*34*). Cultures of *B. subtilis* strains were inoculated in LB broth supplemented with antibiotics as needed and grown overnight in a shaking incubator at 37 ºC and 250 rpm. The overnight cultures were subcultured into fresh LB in 96-deep-well plates and grown for 7h at 37 ºC and 250 rpm. The xylanase activity assays were performed in 96-well clear flat-bottom plates. In each well, 20 µl of culture supernatant and 20 µl of substrate solution (1% beechwood xylan (Neogen, Bray, Ireland) in 10 mM Tris-HCl, pH 8.0) were added. The reactions were carried out at 40°C for 1h. Then, 160 µl of dinitrosalicyclic acid (DNS) solution (0.1% 3,5 dinitrosalicyclic acid, 30% sodium potassium tratrate, pH 7.0) was added to each well, and the microplates were incubated at 100°C for 20 min in a dry oven. Finally, the plates were analyzed using a plate reader (BioTek, Winooski, VT, USA) with Gen 5 software (version 3.03) to determine the optical density of each sample at 570 nm. Standard xylose (cat#X1500 Sigma-Aldrich, St. Louis, MO, USA) solutions were prepared in 10 mM Tris-HCl, pH 8.0, and 40 µl of each standard was analyzed using DNS solution as described above.

**Measurement of pulcherrimin**

Pulcherrimin measurements were performed as previously described (*35*), with some modifications. *B. subtilis* strains were grown overnight in LB broth in a shaking incubator at 37ºC and 250 rpm. The overnight cultures were then subcultured into 500 µl of modified LBGM medium (LB broth supplemented with final concentrations of 0.25% glycerol (v/v), 25 µM $MnSO_4$, and 0.2 mM $FeCl_3$) (*36*)in culture 96-deep-well plates and incubated for 48h at 37ºC and 250 rpm. The cultures were then centrifuged at 3,750 rpm for 10 min at room temperature and washed with 500 µl 1X PBS. The resulting pellets were resuspended in 500 µl of 2M NaOH and incubated for 10 min at room temperature. The cells suspensions were then centrifuged at 3,750 rpm for 10 min at room temperature. Following centrifugation, 200 µl of the supernatant were transferred to the wells of a clear, flat-bottom, 96-well plate and OD at 410 nm was measured using a Synergy H1 plate reader (BioTek, Winooski, VT, USA) with Gen 5 software (version 3.03). OD at 600 nm was measured for each culture and used to normalize the readings.

**Proteomics**

Cells were prepared for proteomic analysis as previously described (*37*, *38*). Briefly, cells were lysed by bead beating with 0.15 mm zirconium oxide beads in 100 mM Tris-HCl, pH 8.0 (Geno/Grinder 2010, SPEX). The crude lysates were adjusted to 4% SDS and 10 mM dithiothreitol, heated to 90°C, and cleared by centrifugation. Protein concentrations were measured by a NanoDrop OneC spectrophotometer (Thermo Scientific, Waltham, MA, USA) at 205 nm. Cleared lysates were moved to a new tube, and cysteines alkylated with 30 mM iodoacetamide. Proteins were isolated via the protein aggregation capture method (*39*), washed as prescribed, and digested in situ with sequencing-grade trypsin at a 1:75 (wt/wt) trypsin-to-protein ratio in 100 mM ammonium bicarbonate, pH 8.0, at 37°C overnight and the following day. The resulting tryptic digests were acidified with formic acid, filtered through a 10 kDa MWCO centrifugal concentrator

(Vivaspin 500, Sartorius, Goettingen, Germany), and quantified by NanoDrop OneC. For each sample, 2 µg of peptides were analyzed by automated 1D LC-MS/MS using a Vanquish ultra-high-pressure liquid chromatography (UHPLC) system connected directly to a nanoelectrospray source on a Q Exactive Plus mass spectrometer (Thermo Scientific, Waltham, MA, USA), as described previously (*38*).

MS/MS spectra were searched against the *B. subtilus* RO-NN-1 and *B. subtilis* NCIB 3610 reference proteomes from UniProt (along with common protein contaminants) using the SEQUEST HT algorithm in Proteome Discoverer version 2.5 (Thermo Scientific, Waltham, MA, USA), and peptide spectrum matches (PSMs) were scored and filtered with Percolator, controlling the false-discovery rate to <1% at the PSM and peptide levels (*40*). Peptides were quantified by the chromatographic area under the curve, mapped to their respective proteins, and summed to estimate protein-level abundance. Raw proteins files were exported from Proteome Discoverer. Log2-transformation of protein abundances was performed followed by local regression (LOESS) normalization and mean-centering across the entire dataset in R using scripts from the InfernoRDN software (v1.1.7995) (*41*). The abundance values for proteins with missing values were imputed with random values drawn from the normal distribution (width 0.3, downshift 2.5) using R. For each protein, an ANOVA was run using the package rstatix (*42*)with a Benjamini Hochberg correction for multiple comparisons followed by Tukey pairwise post-hoc tests to test for differences between genotype and growth phase.

**Quantitative analysis of surfactin production**

Cell pellets were flash-frozen and lyophilized until completely dry. A biphasic extraction protocol was performed as previously described (*43*). Briefly, equal volumes of ice-cold hydrated ethyl acetate and LC-MS grade water were added to the samples, using 200 µL for cell pellets. Samples were vortexed for 1 minute and incubated overnight at 4°C. Following a 10-minute centrifugation, the ethyl acetate and aqueous fractions were separated. The aqueous fractions were then filtered using a 10 kDa MWCO centrifugal concentrator (Vivaspin 500, Sartorius, Goettingen, Germany) by centrifugation at 12,000 × g. The filtered aqueous extracts were freeze-dried and resuspended in an aqueous solvent (5% acetonitrile, 0.1% formic acid), using 30 µL for cell pellets. The ethyl acetate extracts were air-dried in a chemical fume hood and subsequently resuspended in an organic solvent (70% acetonitrile, 0.1% formic acid) using the same volumes as the aqueous fractions.

Following resuspension, both aqueous and organic extracts were analyzed using untargeted metabolomics via UHPLC-MS/MS to profile metabolite composition. Samples were analyzed using a ThermoFisher Q-Exactive Plus mass spectrometer equipped with an in-house-constructed nanospray analytical column (75 µm × 150 mm) packed with a 1.7-µm C18 Kinetex RP C18 resin (Phenomenex). The mobile phase consisted of solvent A (95% water, 5% acetonitrile, 0.1% formic acid) and solvent B (70% acetonitrile, 30% water, 0.1% formic acid). Metabolite separation was achieved using a 30-minute linear gradient from 5% aqueous solvent to 100% organic solvent at a flow rate of 250 nL/min, with an injection volume of 10 µL followed by mass spectrometric detection in positive electrospray ionization (ESI+) mode. Full-scan MS spectra were acquired using Xcalibur v4.3 from 135 to 2000 m/z at 70,000 resolution by employing the top N data-dependent acquisition method, where N is 5. Fragmentation of precursor ions was performed using stepped higher-energy C-trap dissociation at collision energies of 10, 20, and 40 eV. To prevent redundant sampling of abundant metabolites, a dynamic exclusion of 10 seconds was applied.

Untargeted LC-MS/MS data were processed using Skyline v24.1.0 (*44*, *45*) to identify and quantify select surfactin compounds. Raw spectral data were imported into Skyline, where surfactins were detected based on specified m/z ratios and corresponding fragmented ions. Retention time alignment was performed across all data files to maintain consistency in peak detection and improve comparability between samples. Processed LC-MS/MS data were then further analyzed using R v4.4.1, with surfactin log10-transformed for normalization before statistical analysis. Statistical comparisons were conducted in R using the ggpubr package (cite). Pairwise comparisons between strains were conducted using Student's t-test, with Bonferroni correction applied to adjust for multiple comparisons. P-values were annotated directly on the boxplots to indicate statistical significance, with asterisks representing different levels of significance.

**Strain construction**

*B. subtilis* DK1042 HK (JMB92) and DK1042 ME (JMB90) strains were generated by amplifying allele replacement constructs from genomic DNA of strains BKK34900 (168 Δ*hisB*::*kan*) and BKE13180 (168 Δ*metE*::*erm*), respectively, and transforming strain DK1042 with these gene targeting constructs via natural competency. Transformants were selected on LB agar supplemented with kanamycin and erythromycin plus lincomycin, respectively.

*B. subtilis* DK1042 HK ME (JMB91) was constructed by transforming strain DK1042 HK with a *metE*::*erm* allele replacement construct via natural competency. Transformant were selected on LB agar supplemented with kanamycin, erythromycin and lincomycin.

*B. subtilis* NCIB 3610 ΔpBS32 (JMB265) was generated by transforming *B. subtilis* DK1042 with pJM493 via natural competency. Transformants were selected on LB agar plates supplemented with kanamycin and 200 ng/ml tetracycline at 30 ºC. Single colonies were then streaked on LB plates and incubated at 45 ºC overnight to cure pJM493. Absence of pBS32 was validated by whole-genome resequencing.

*B. subtilis* RO-NN-1 *gerA*::*kanR* (JMB266) was constructed by transforming *B. subtilis* RO-NN-1 with a DNA fragment generated by splicing by overlap extension (SOE) PCR of three PCR products: (1) 547 bp fragment upstream of *gerAA* amplified with primers and genomic DNA from strain RO-NN-1 as template, (2) 562 bp fragment downstream of *gerAC* amplified with primers and genomic DNA from strain RO-NN-1 as a template, and (3) kanamycin resistance cassette amplified with primers and genomic DNA from strain BKK34900 (*6*) as template.

Strains JMB267, JMB272 and JMB274 were constructed by cotransforming *B. subtilis* DK1042 with pJM494 and a repair template generated by amplification of the *gerAA* region from genomic DNA of strain RO-NN-1. Transformants were selected on LB agar plates supplemented with kanamycin and 200 ng/ml tetracycline at 30 ºC. Single colonies were then streaked on LB agar plates and incubated at 45 ºC overnight to cure pJM494.

Strains JMB352-JMB374 were generated by transforming *B. subtilis* P1A7, P1A11, P1B5, P1B12, P1E9, P1F11, P2B1, P2D3, P2D4, P2E11, P2G8, P2G10, P3B5, P3C6, P3D3, P3E3, P3F1, P3F10, P3G3, P3G12, P3H8, P4A9, and P4B5 with pDR244 (*6*), a temperature-sensitive plasmid expressing the Cre recombinase. Transformants were selected on LB agar plates supplemented with spectinomycin at 30 ºC. Single colonies were then streaked on LB agar plates and LB agar plates supplemented with kanamycin (for HK strains) or erythromycin (for ME strains). Strains that grew on LB agar plates, but not on LB agar plates supplemented with antibiotics were streaked on LB agar plates and grown overnight at 45 ºC to cure pDR244. Single transformants were patched onto LB agar plates and LB agar plates supplemented with spectinomycin. Select strains that did no longer grow on spectinomycin plates were validated by whole-genome resequencing.

### Supplementary Text

Subhead

Type or paste text here. This should be additional explanatory text, such as: extended technical descriptions of results, full details of mathematical models, extended lists of acknowledgments, etc. It should not be additional discussion, analysis, interpretation, or critique.

First, we replaced amino acid biosynthesis genes, located roughly in the opposite regions of the chromosome, with antibiotic resistance markers in RO-NN-1 and DK1042, and then crossed the resulting auxotrophic strains with wild-type DK1042 and RO-NN-1, respectively. In each cross, we selected for recombinant strains containing all possible combinations of markers from both parents, generating S1 progeny. A pool of S1 strains was then backcrossed to strain DK1042 creating S2 progeny and the same process was repeated once more to generate S3 progeny. We isolated and sequenced a set of recombinant strains from the S2 and S3 shuffles to evaluate the density of recombination events across their genomes. Recursive protoplast fusion efficiently increased genome-wide recombination with strains from the S3 progeny inheriting, on average, 17.0% of genomic fragments from strain DK1042 creating extensive genetic diversity across their chromosomes that could be used for genetic mapping. Although no reference data are available to guide us on the minimum amount of genome-wide recombination necessary for efficient genetic mapping in bacteria, this level of genetic mixing surpasses that typically achieved in traditional eukaryotic biparental crosses. Successive rounds of shuffling and selection introduced long recombination fragments originating from strain DK1042 flanking the selected markers. To introgress these regions, we generated a set of S4 recombinant strains by backcrossing a pool of S3 strains to strain RO-NN-1. We determined the genome architecture of 573 individual S3 strains and 50 S4 strains. After removing nearly identical strains and quality control, we retained a total of 350 shuffled strains with 60,937 SNPs (minor allele frequency of at least 0.03) as a mapping population.

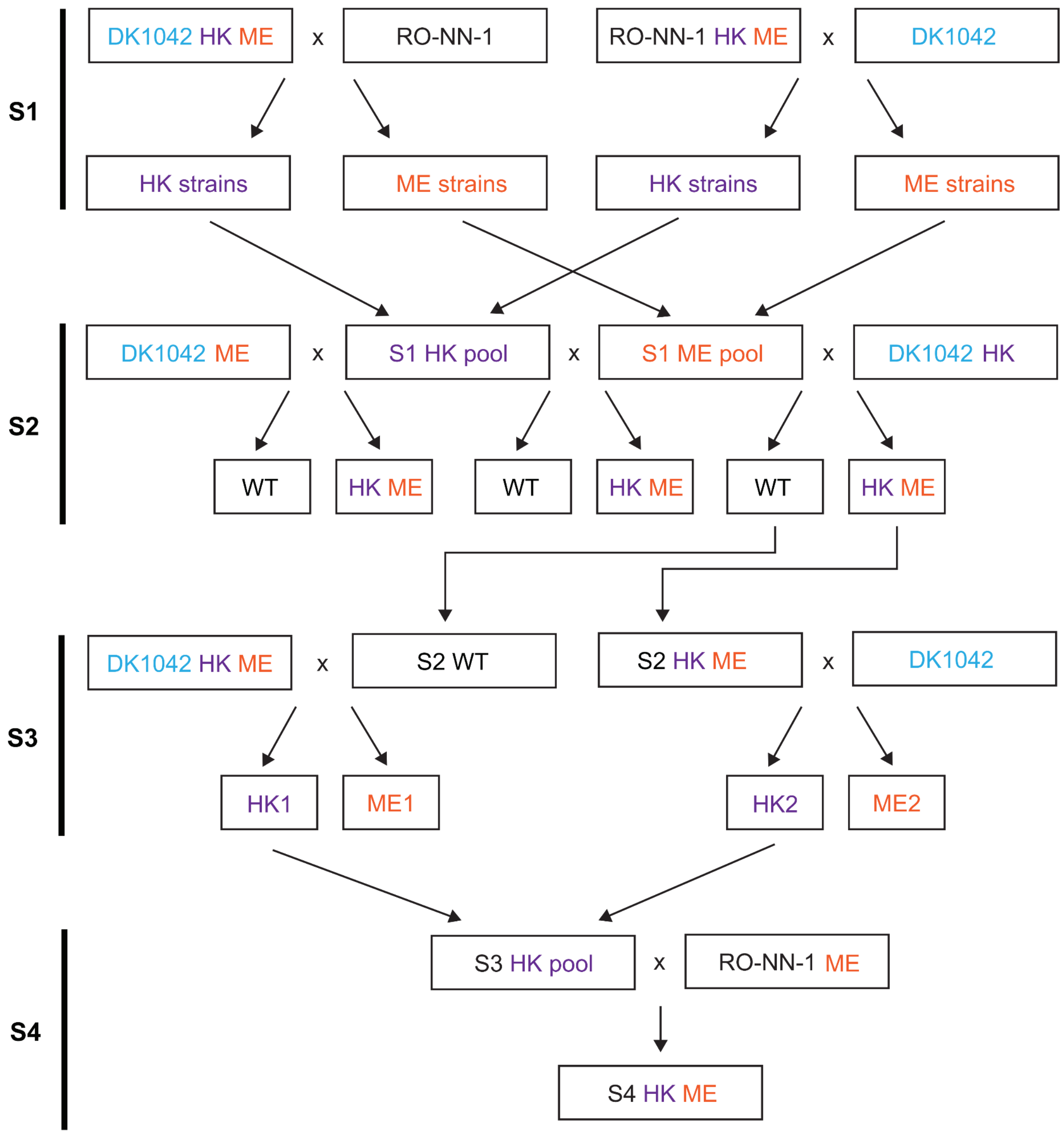

**Fig. S1. Schematic overview of the construction of the *B. subtilis* QTL mapping population.** A mapping population of recombinant strains was generated through three and four rounds of genome shuffling via protoplast fusion, using *B. subtilis* RO-NN-1 and DK1042 as parental strains, as described in the Materials and Methods section. S1, S2, S3, and S4 indicate the shuffling rounds. WT: wild-type; HK: Δ*hisB*::*kan*; ME: Δ*metE*::*erm*.

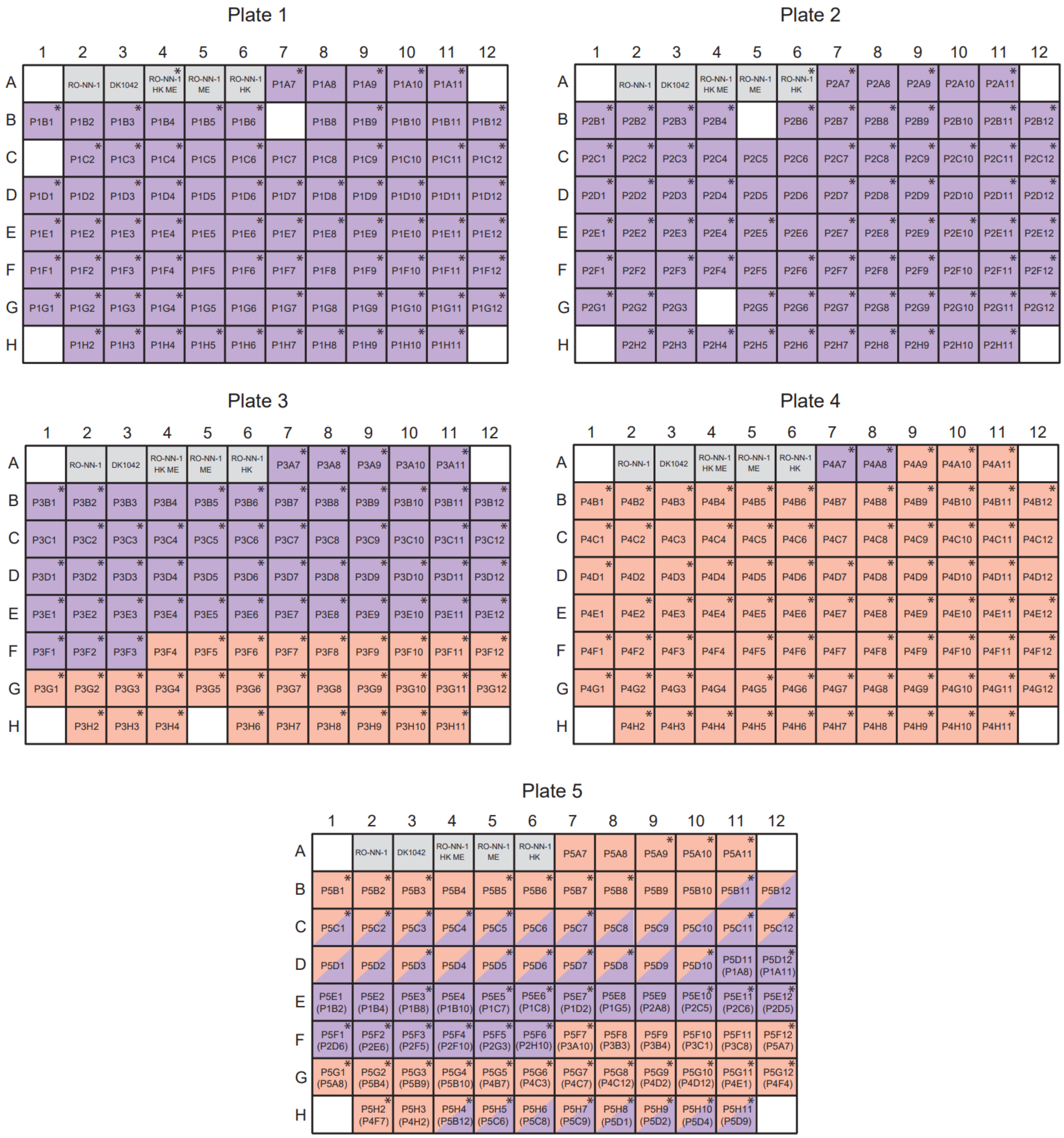

**Fig. S2. Layout of the 96-well plates with *B. subtilis* shuffled strains generated in this study.** Wells containing HK and ME shuffled strains are highlighted in purple and orange, respectively. Wells containing HK ME double-resistant shuffled strains are highlighted using both purple and orange. Blank wells are empty. *B. subtilis* RO-NN-1, DK1042, RO-NN-1 HK ME, RO-NN-1 HK and RO-NN-1 ME are included on each plate, with their corresponding wells highlighted in gray. Plate 5 includes randomly selected replicates, indicated in parentheses. The 350 genetically diverse strains used for genetic mapping are marked with asterisks. The five plates have been deposited to BGSC. HK: Δ*hisB*::*kan*; ME: Δ*metE*::*erm.*

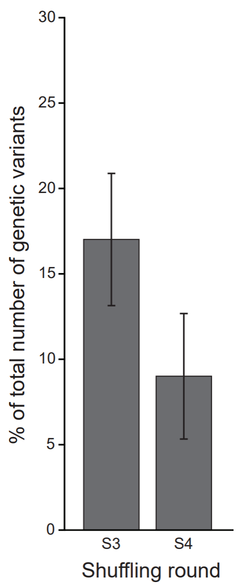

**Fig. S3. Genome-wide recombination in the *B. subtilis* QTL population.**
The fraction of the genome that was recombined after the S3 and S4 shuffling rounds between strains RO-NN-1 and DK1042 is shown. S3 and S4 shuffled strains were generated as described in the Materials and Methods and fig. S1. Data represent the mean results from 24 randomly selected S3 strains and all 24 S4 strains. Error bars indicate standard deviation.

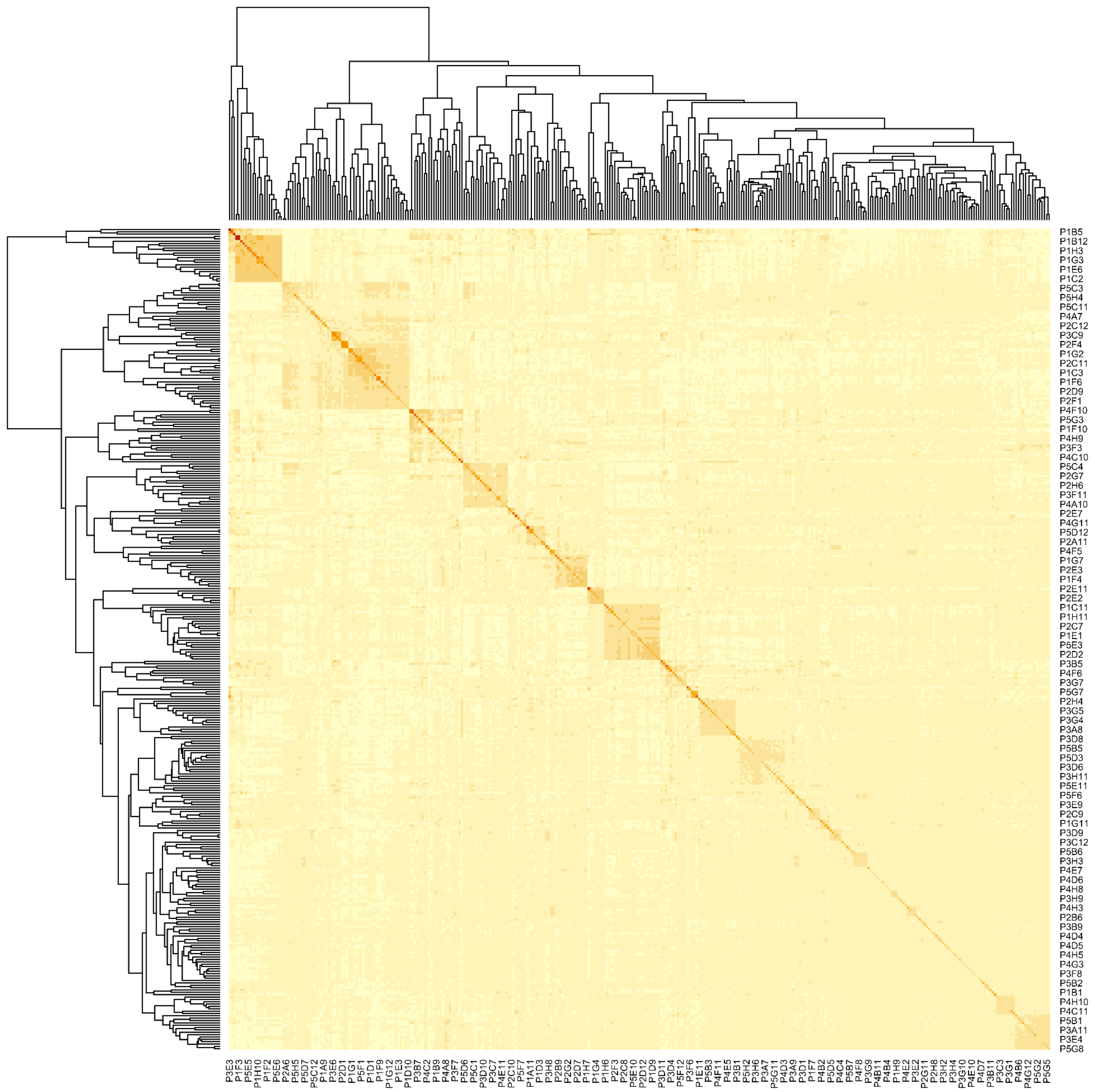

**Fig. S4. Heatmap and dendrogram of a genetic relationship matrix of the *B. subtilis* QTL population.**

The kinship matrix was estimated using the Zhang algorithm (*22*) based on 60,937 SNPs among 350 shuffled strains.

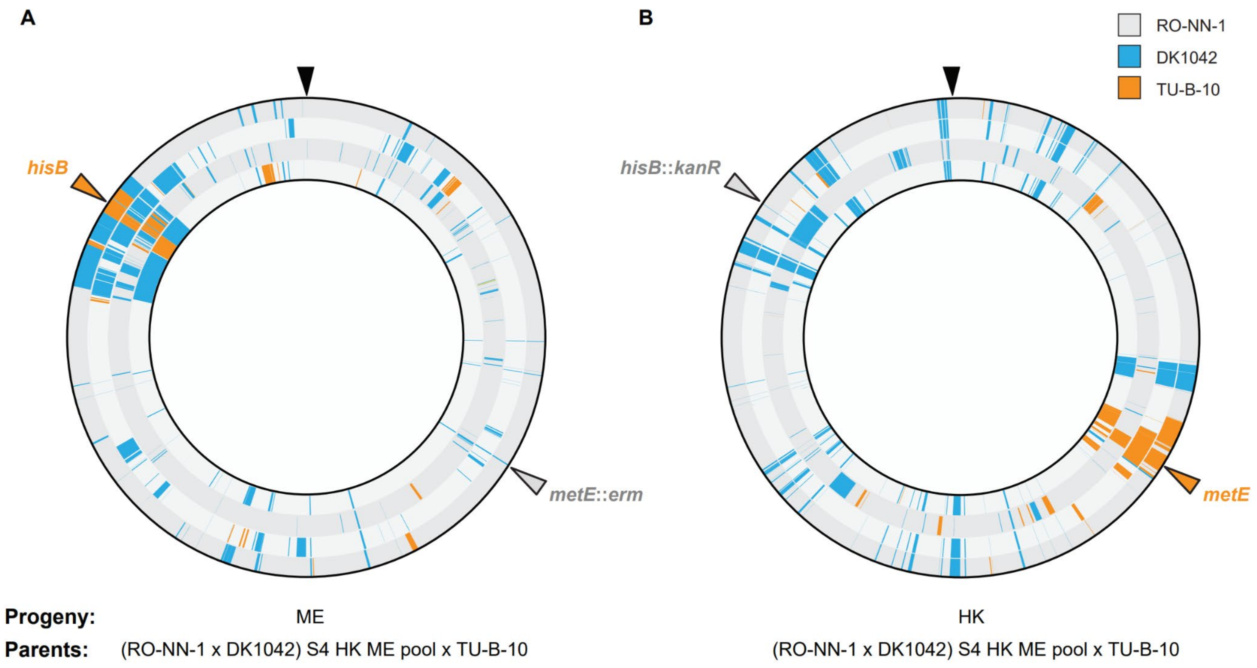

**Fig. S5. Multiparental genome shuffling can maximize allelic diversity in bacterial QTL mapping populations.**
Crossing the *B. subtilis* S4 HK ME pool (Fig. S1) and *B. spizizenii* TU-B-10 generated (**A**) ME and (**B**) HK progeny. Each concentric circle represents an individual sequenced shuffled strain from that cross. Blue bars indicate recombination fragments originating from DK1042. Orange bars represent sequences recombined from strain TU-B-10. The remaining genomic sequences in grey come from strain RO-NN-1. Orange and grey arrows indicate locations of the selection markers. Black arrows indicate the origin of replication. ME: Δ*metE*::*erm*; HK: Δ*hisB*::*kan*.

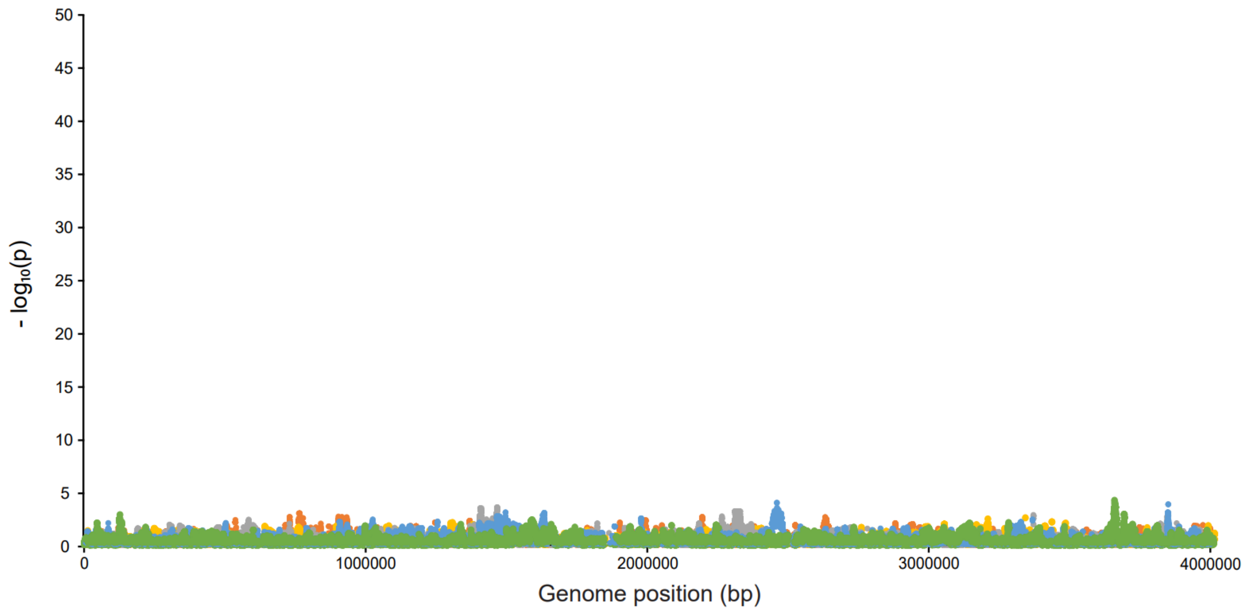

**Fig. S6. Manhattan plot using randomized spore germination phenotypic data demonstrates no significant association in the *B. subtilis* QTL population.**
Intrinsic noise from genetic variation was estimated by running QTL mapping permutations with real genotypic data and randomized binary phenotypes. The colors indicate association results from different permutations.

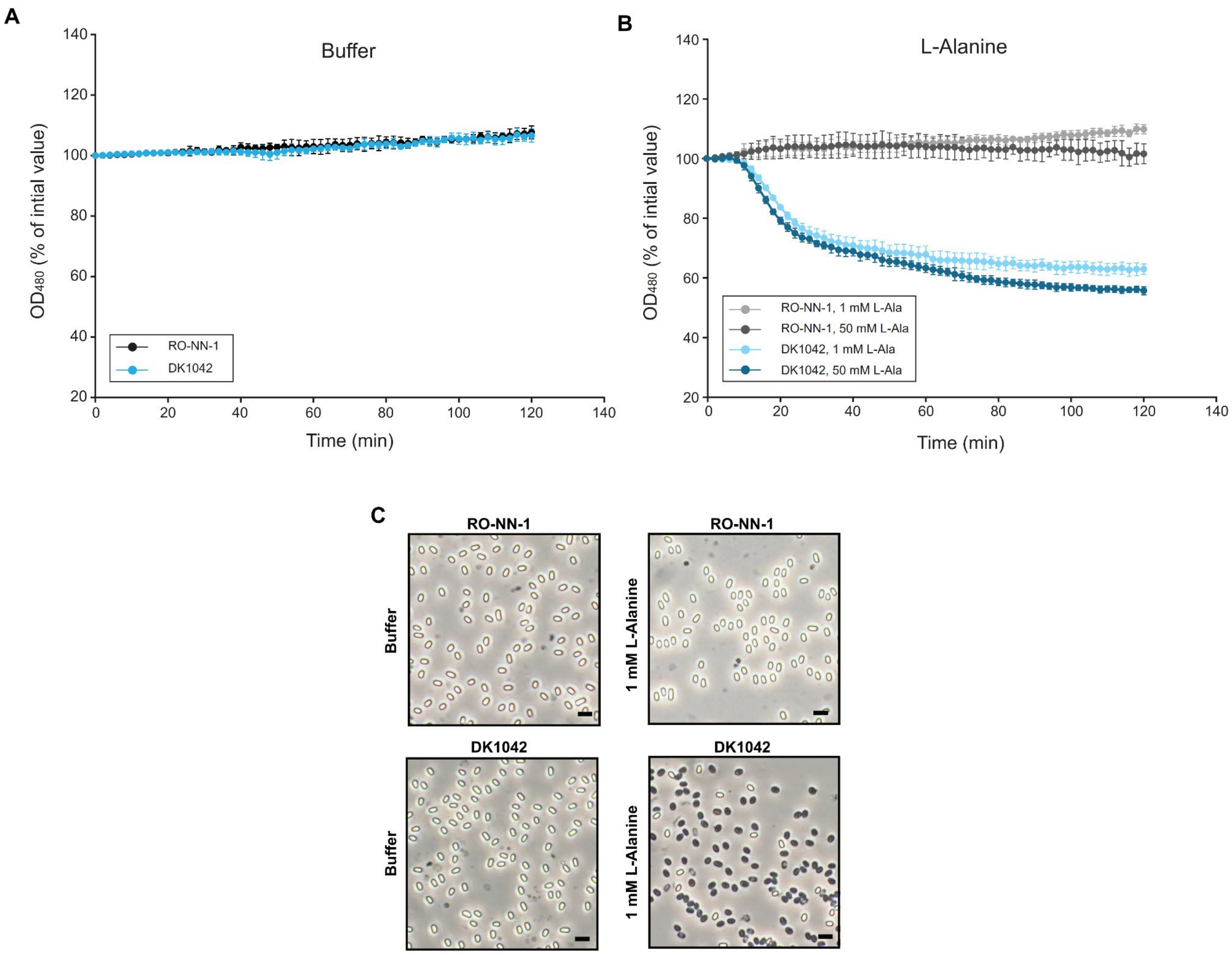

**Fig. S7. Spore germination of *B. subtilis* RO-NN-1 and DK1042 in response to L-alanine.** Spores of strains RO-NN-1 and DK1042 were incubated with (**A**) buffer and (**B**) 1 mM or 50 mM L-alanine, and spore germination was assessed as percent reduction in $OD_{480}$. (**C**) Representative phase-contrast images of RO-NN-1 and DK1042 spores after 120 min incubation with buffer and 1 mM L-alanine. Data represents the mean results of three biological replicates. Error bars indicate standard deviation. Scale bar, 2 μm.

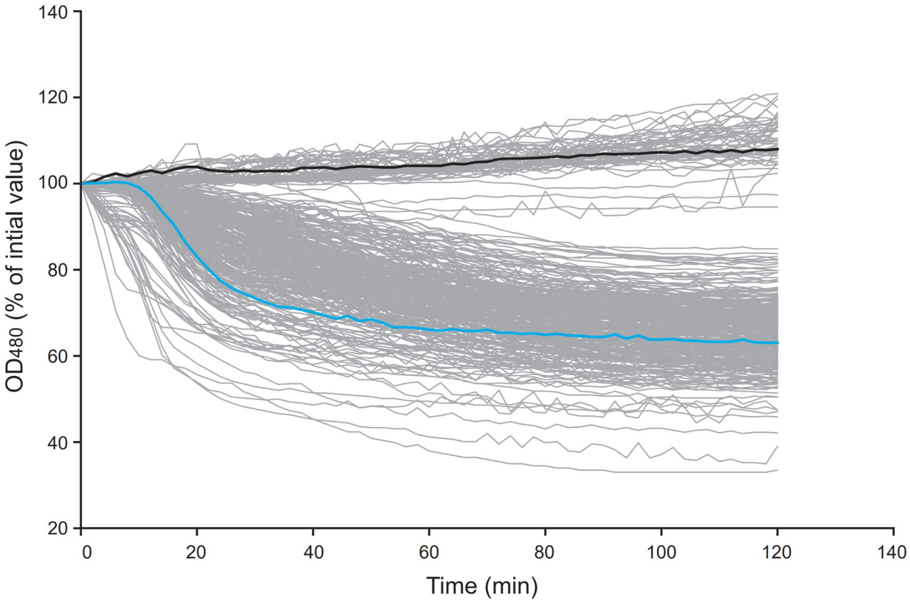

**Fig. S8. Spore germination of the *B. subtilis* QTL population in response to 1 mM L-alanine.**

Spores of the 350 shuffled strains and the QTL parents were incubated with 1 mM L-alanine, and spore germination was assessed as percent reduction in initial $OD_{480}$ over the course of 120 min. Gray, black and blue lines indicate data from the 350 shuffled strains, RO-NN-1 and DK1042, respectively. Data from one biological experiment is shown.

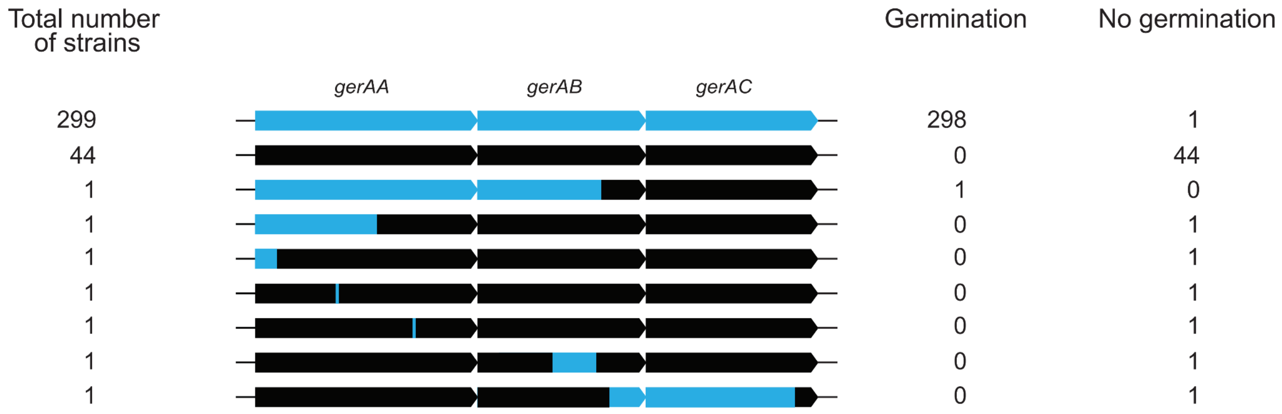

**Fig. S9. Distribution of candidate causative spore germination loci in the *B. subtilis* QTL mapping population.**

Blue and black regions indicate sequences originating from *B. subtilis* DK1042 and RO-NN-1, respectively. The number of strains from each genotype and their respective phenotypes are shown.

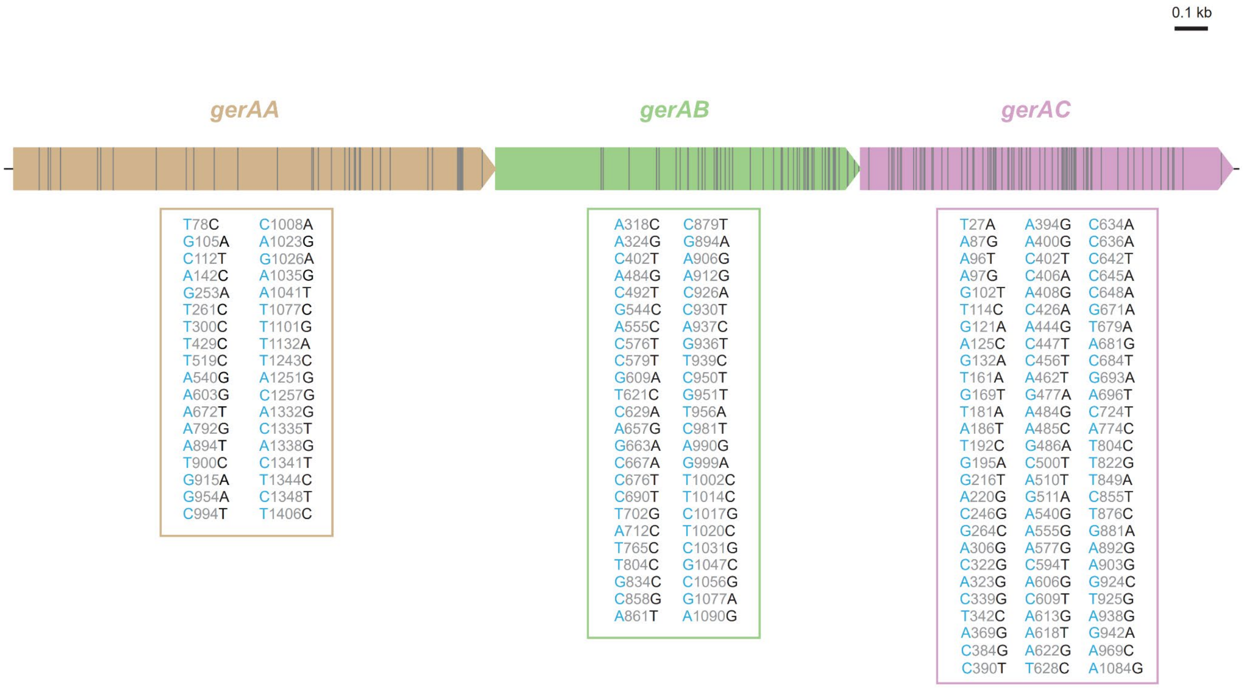

**Fig. S10. Overview of genetic variants within the *gerA* locus.**
Blue and black letters indicate SNPs in DK1042 and RO-NN-1, respectively.

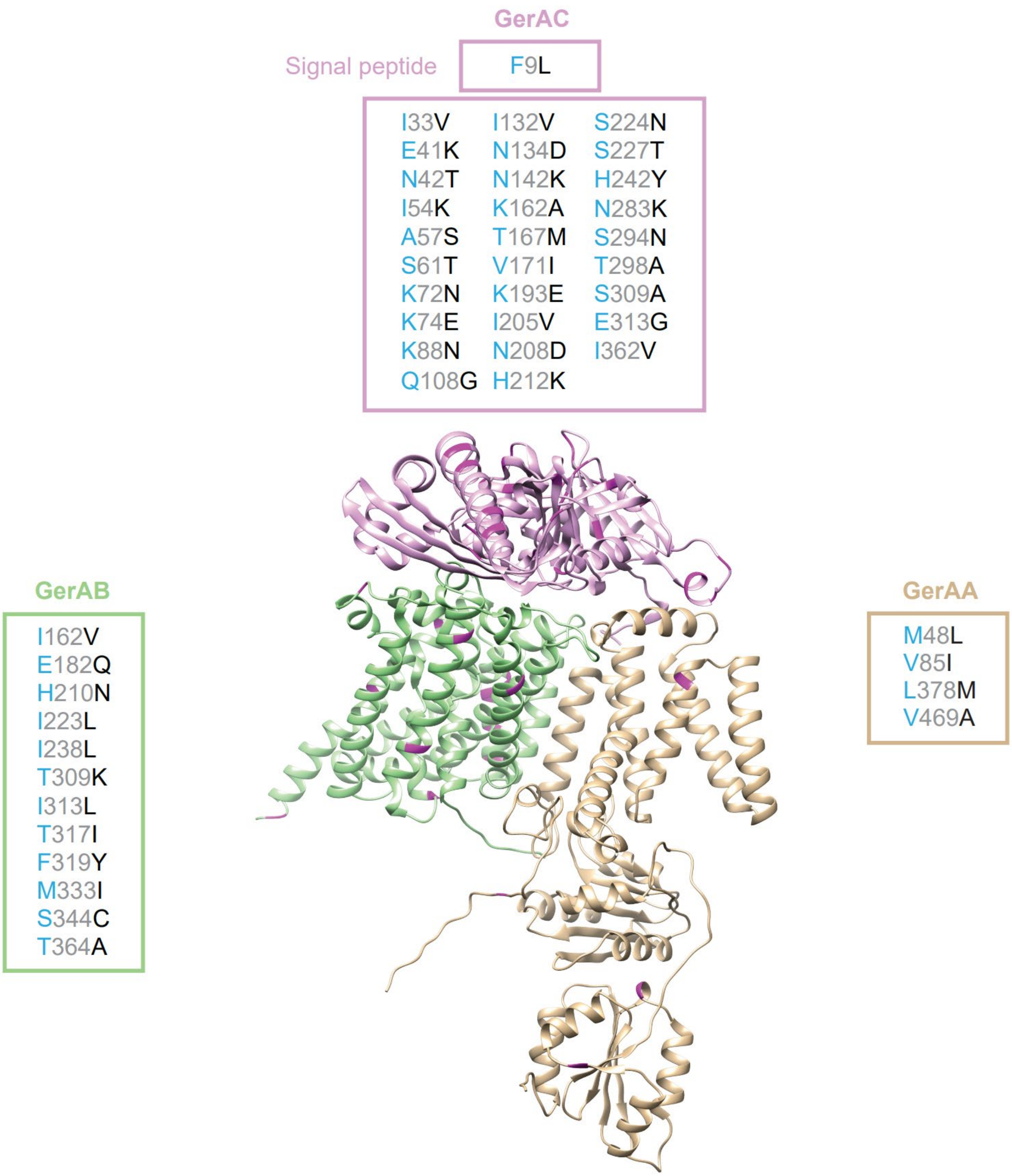

**Fig. S11. Overview of amino acid differences in the GerAA, GerAB, and GerAC proteins between *B. subtilis* RO-NN-1 and DK1042.**

Blue and black letters indicate SNPs in DK1042 and RO-NN-1, respectively.

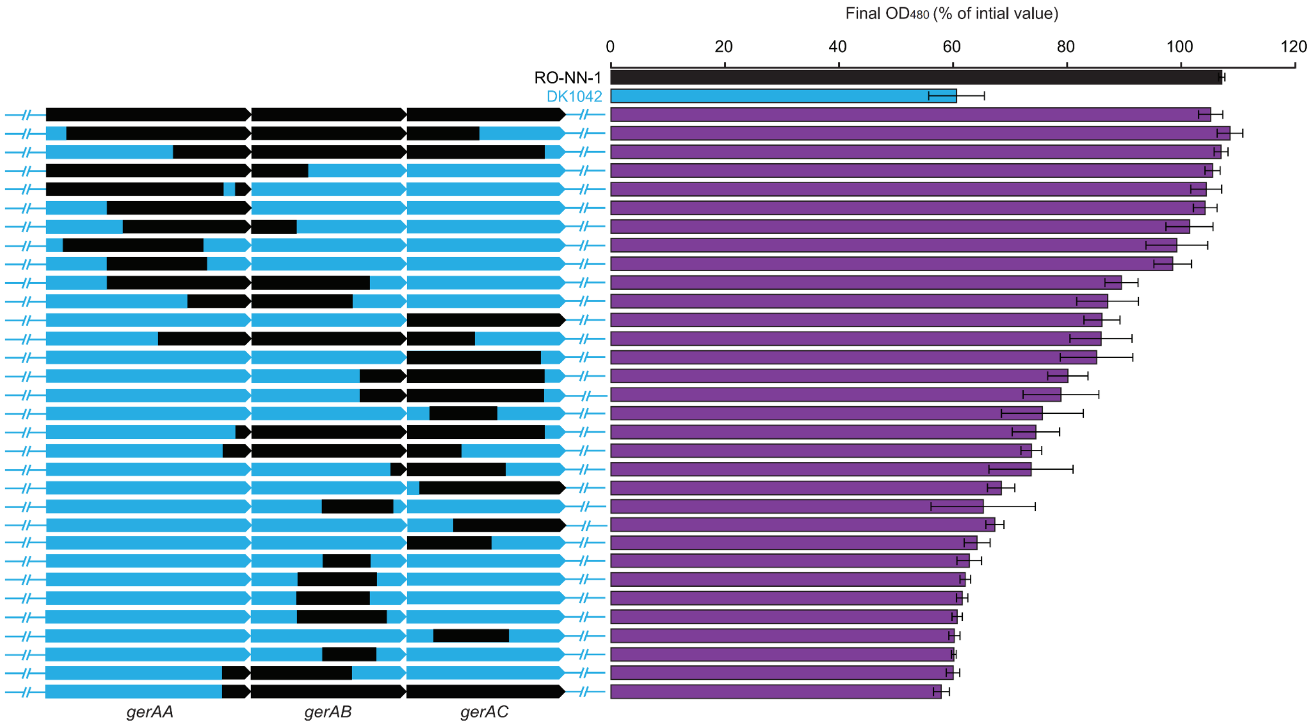

**Fig. S12. Genetic swaps validate causal genetic loci for spore germination in response to L-Alanine.**

Gene segments within the *gerA* operon in strain DK1042 were replaced with corresponding segments from strain RO-NN-1 using a CRISPR-based approach and their spore germination abilities were evaluated. The bar plot shows optical densities measured after incubation of spores of each strain with 1 mM L-Alanine for 120 min. Data are presented as a percentage of the initial $OD_{480}$ of the phase-bright spores. Blue and black regions indicate sequences originating from *B. subtilis* DK1042 and RO-NN-1, respectively.

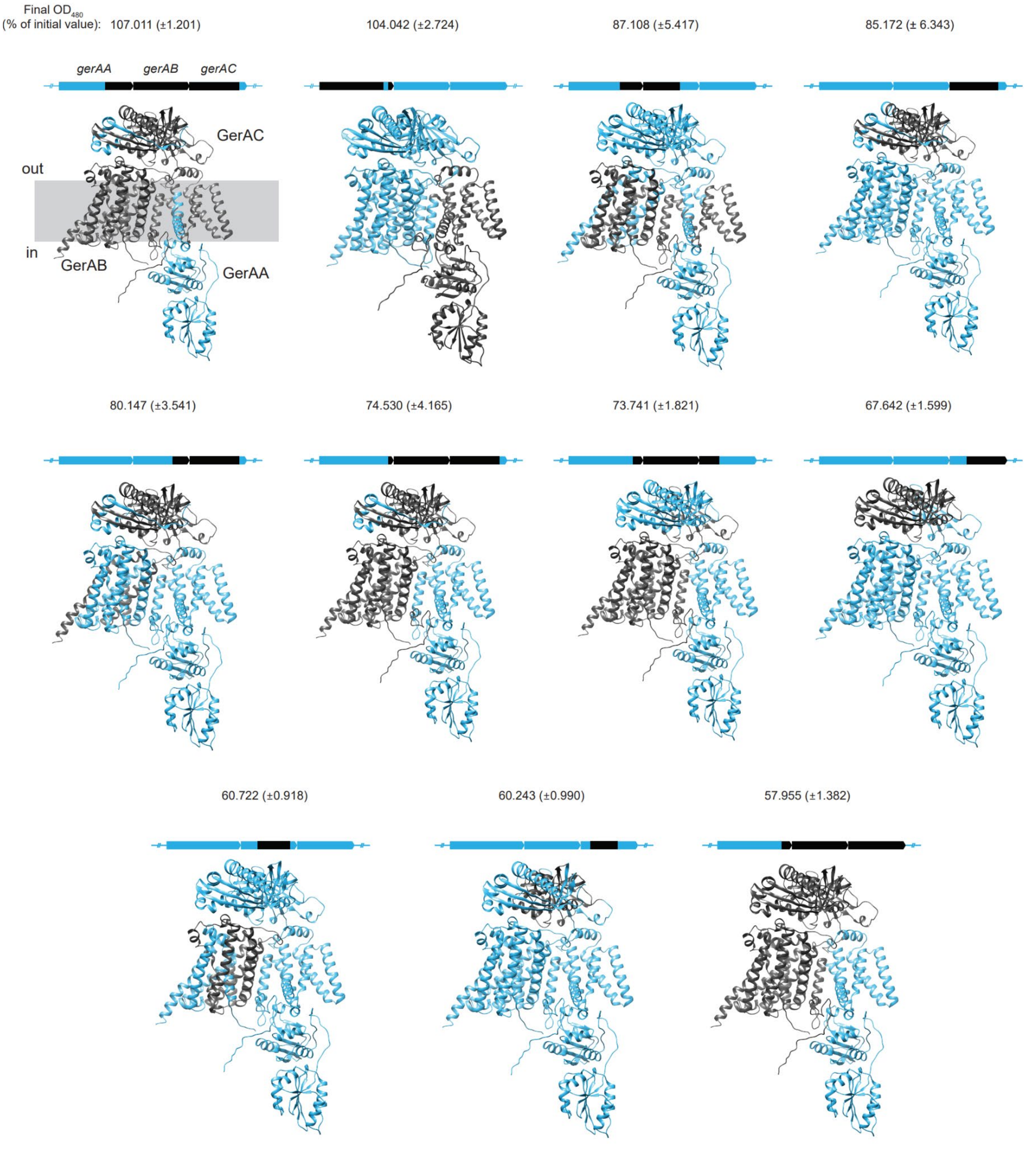

**Fig. S13. Structural models of the GerA trimers resulting from genetic swaps in the *gerA* locus shown in Fig. 2D.**

Final $OD_{480}$ values presented as a percentage of the initial $OD_{480}$ of the phase-bright spores are indicated. The data represents the mean results of three biological replicates, with standard deviations shown in parentheses. Blue and black regions indicate sequences originating from *B. subtilis* DK1042 and RO-NN-1, respectively.

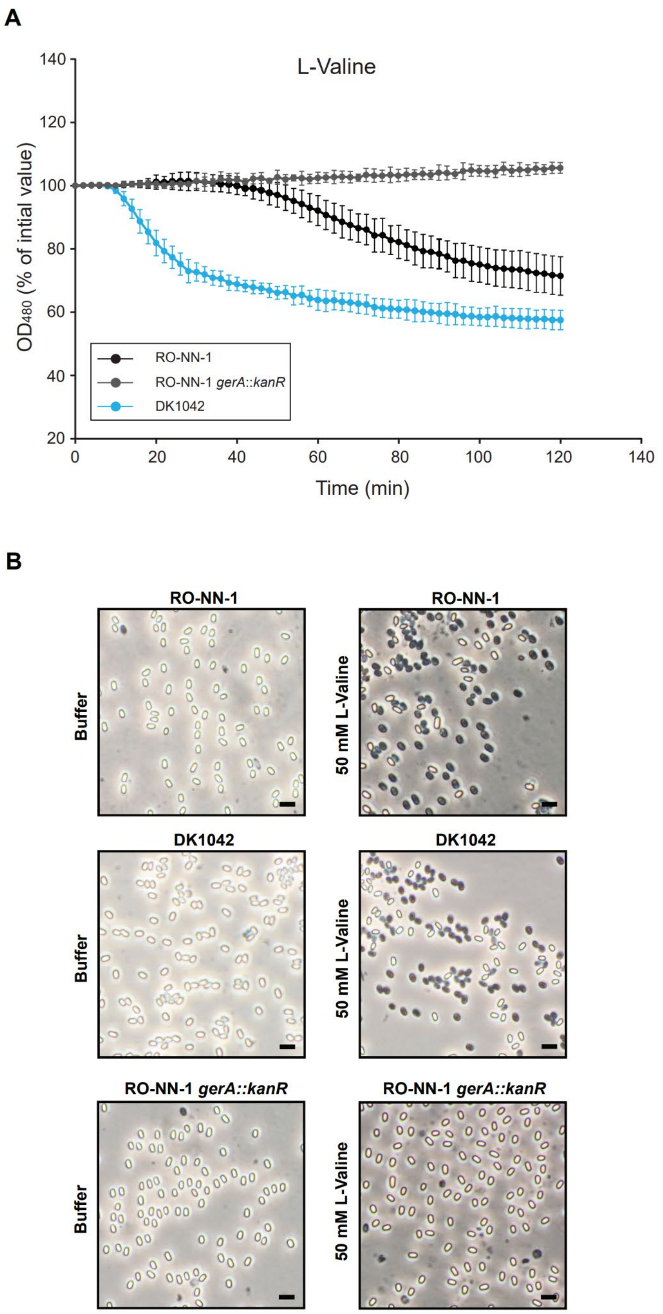

**Fig. S14. Spores of *B. subtilis* RO-NN-1 and DK1042 germinate in response to L-valine.**
(**A**) Spores of strains RO-NN-1, RO-NN-1 *gerA*::*kanR* and DK1042 were incubated with 50 mM L-valine, and spore germination was assessed as percent reduction in $OD_{480}$. (**B**) Representative phase-contrast images of RO-NN-1, RO-NN-1 *gerA*::*kanR* and DK1042 spores after 120 min incubation with buffer and 1 mM L-valine. The data represents the mean results of three biological replicates. Error bars represent standard deviation. Scale bar, 2 μm.

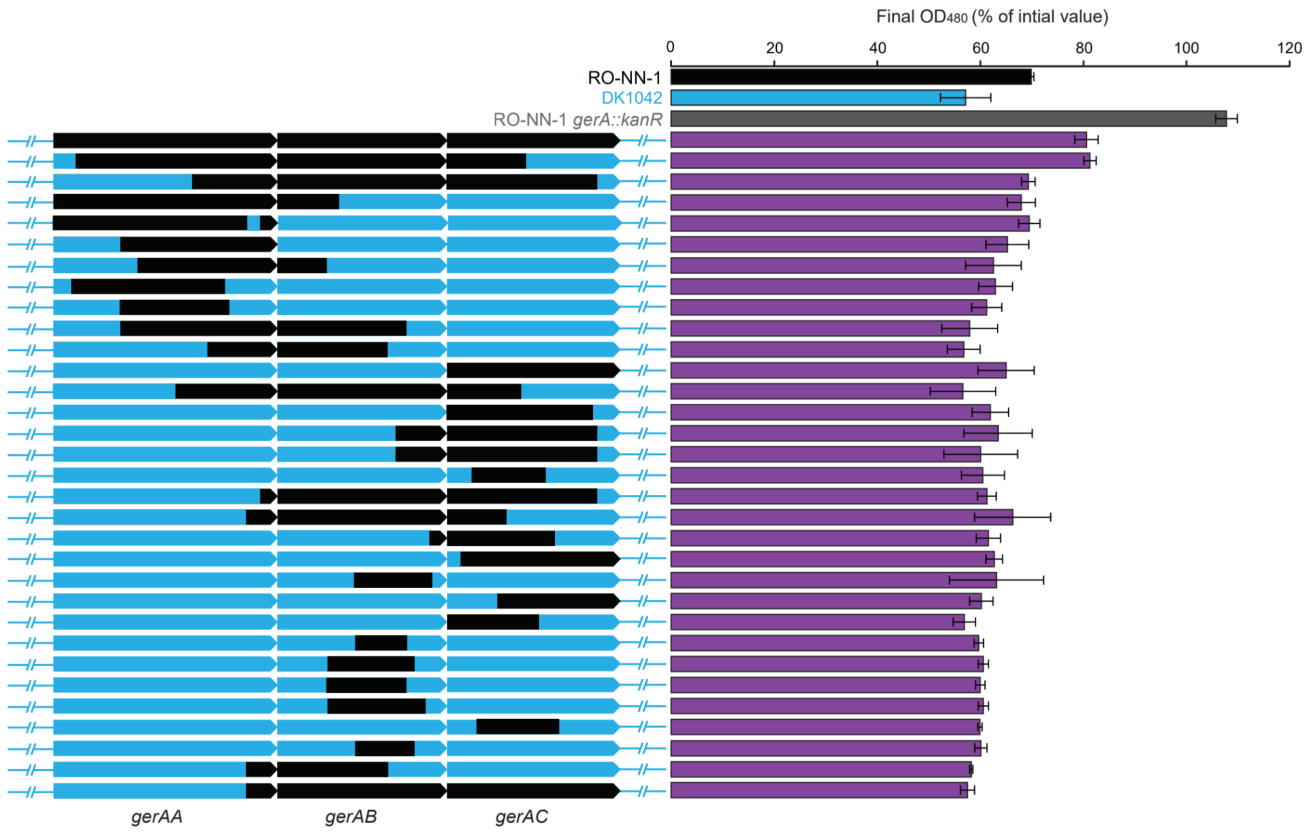

**Fig. S15. Spore germination of strains with genetic swaps within the *gerA* locus in response to L-Valine.**

Gene segments within the *gerA* operon in strain DK1042 were replaced with corresponding segments from strain RO-NN-1 using a CRISPR-based approach and their spore germination abilities were evaluated. The bar plot shows optical densities measured after incubation of spores of each strain with 50 mM L-Valine for 120 min. Data are presented as a percentage of the initial $OD_{480}$ of the phase-bright spores. Blue and black regions indicate sequences originating from *B. subtilis* DK1042 and RO-NN-1, respectively.

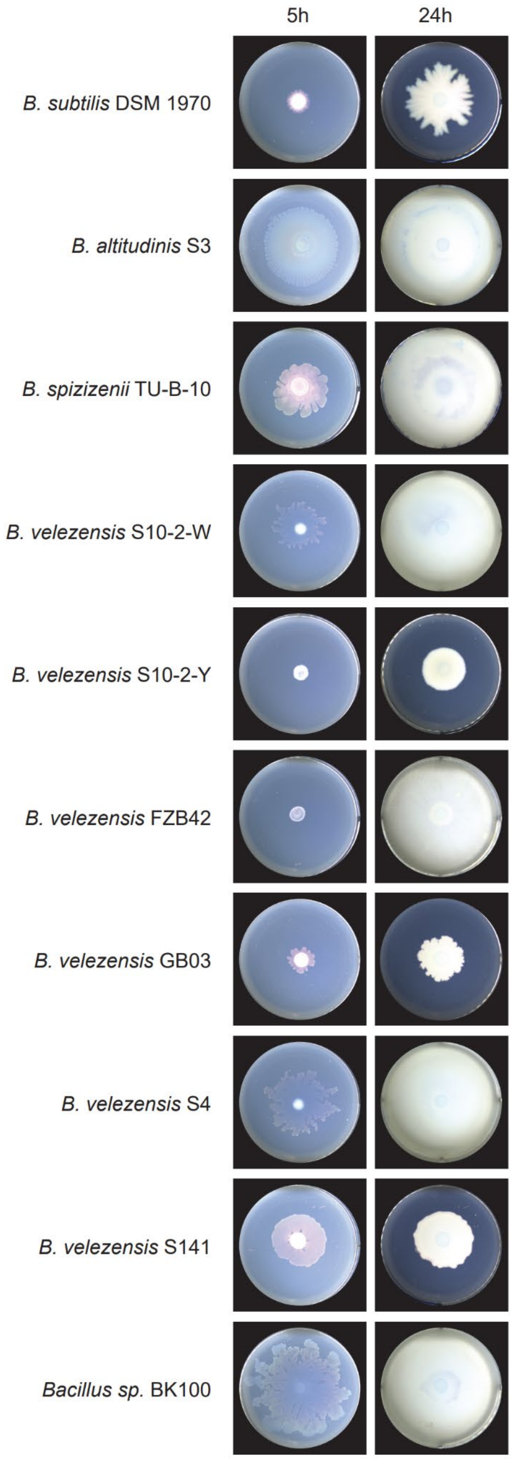

**Fig. S16. Swarming motility varies among different *Bacillus* isolates.**
Swarming motility of *Bacillus* isolates was tested on LB medium solidified with 0.7% agar for 5h and 24h. Representative images from three independent biological experiments are shown.

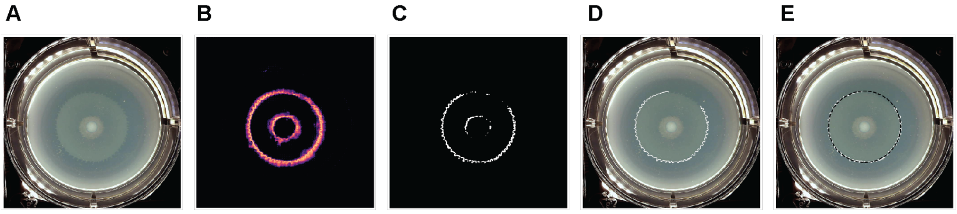

**Fig. S17. Swarm image analysis workflow.**
(**A**) A convolutional neural network (CNN) inputs an image of a *B. subtilis* swarm in a single well of a 6-well plate using the XY-robot. (**B**) The CNN outputs pixel-level probabilities for swarm boundary classification. **(C)** A global threshold is applied to the probability values to create a binary mask. (**D**) The longest continuous segment of the thresholded mask is selected for circle fitting. (**E**) The best fit circle is computed based on the longest segment and subsequently used to estimate swarming phenotypes.

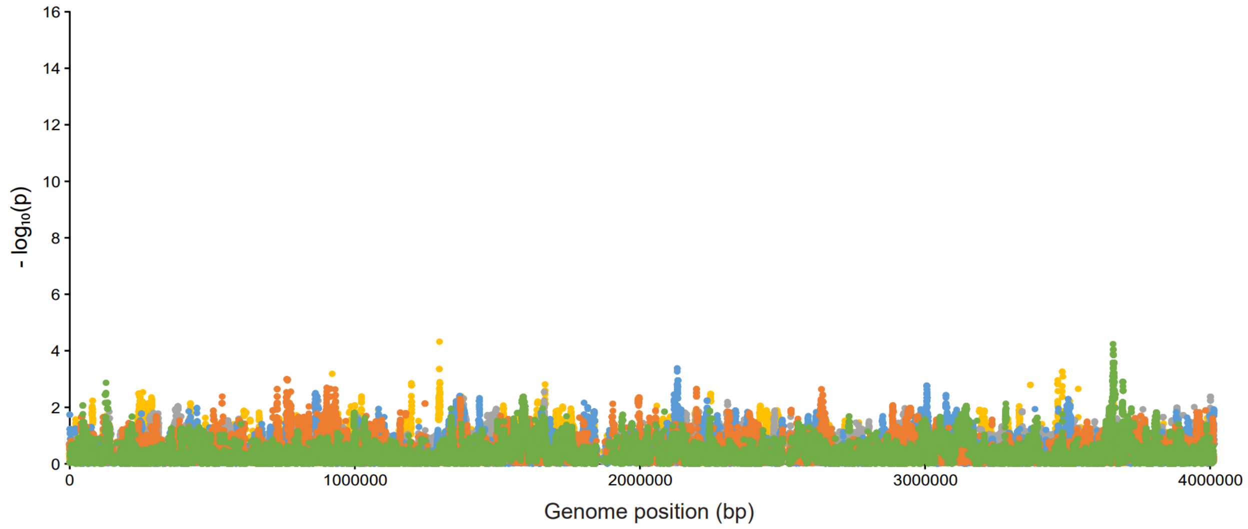

**Fig. S18 Manhattan plot using randomized swarming motility phenotypic data demonstrates no significant association in the *B. subtilis* QTL population.**

Intrinsic noise from genetic variation was estimated by running QTL mapping permutations with real genotypic data and randomized quantitative phenotypes. The colors indicate association results from different permutations.

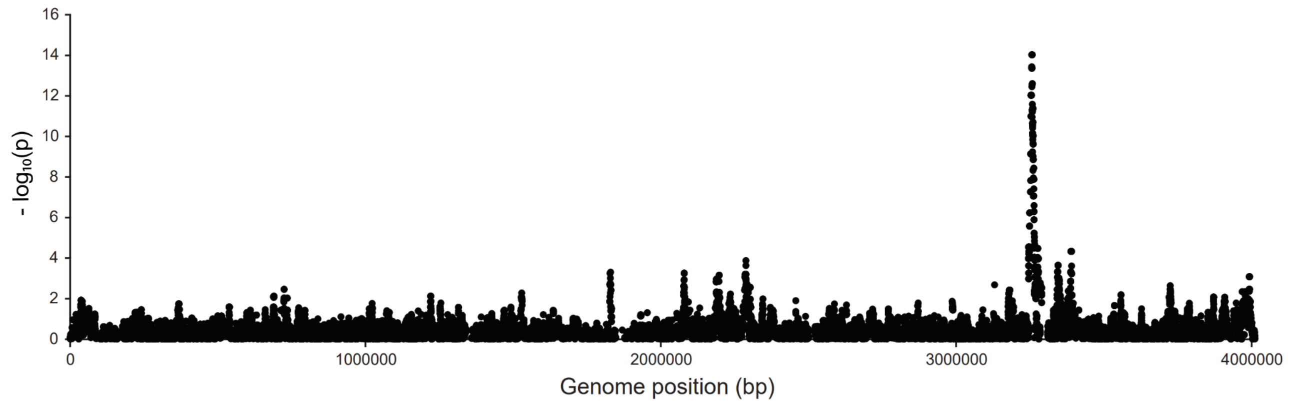

**Fig. S19. Bacterial QTL mapping identifies a highly significant genomic region associated with swarming motility.**

Manhattan plot summarizing the statistical significance of associations between each genetic variant along the genome and swarming motility across the *B. subtilis* QTL mapping population. An inset showing a 50-kb genomic region including the QTL for swarming motility is presented in Fig. 3D.

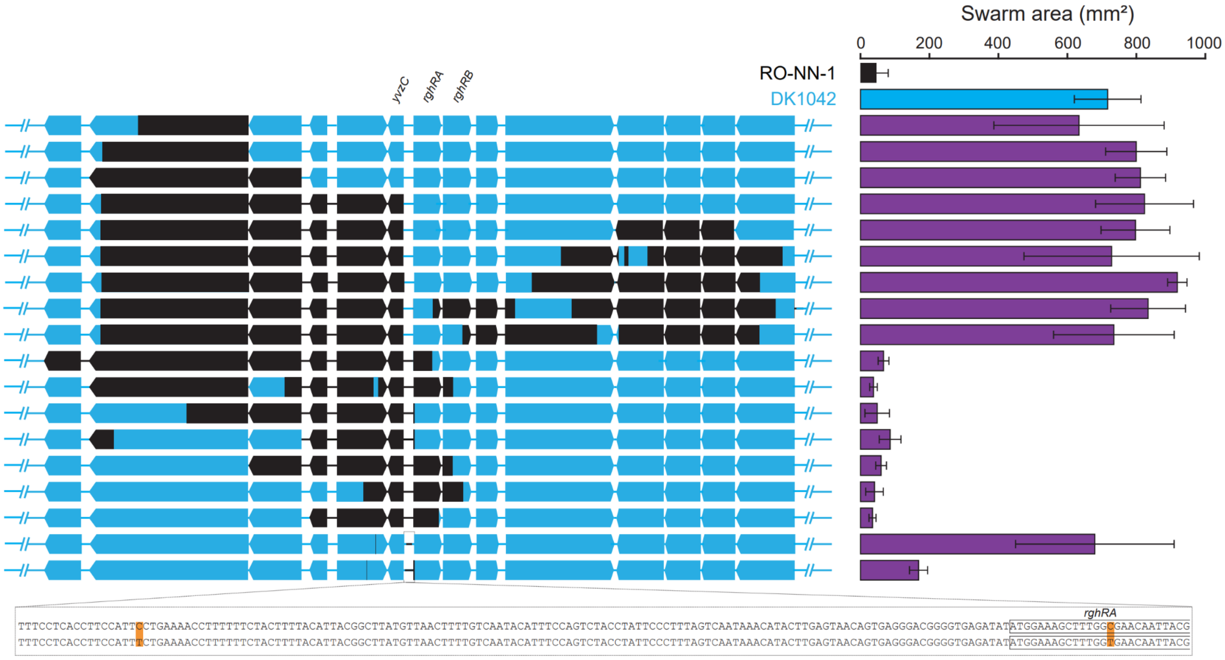

**Fig. S20. Genetic swaps validate causal genetic loci for swarming motility.**
Gene segments within the region *smpB*-*opuBA* in strain DK1042 were replaced with corresponding segments from strain RO-NN-1 using a CRISPR-based approach and the swarming motility areas of the resulting strains were measured. The bar plot shows swarm areas measured after incubation for 5h. The DNA sequence of the genome region containing the two candidate causal genetic variants highlighted in orange is shown. Blue and black regions indicate sequences originating from *B. subtilis* DK1042 and RO-NN-1, respectively.

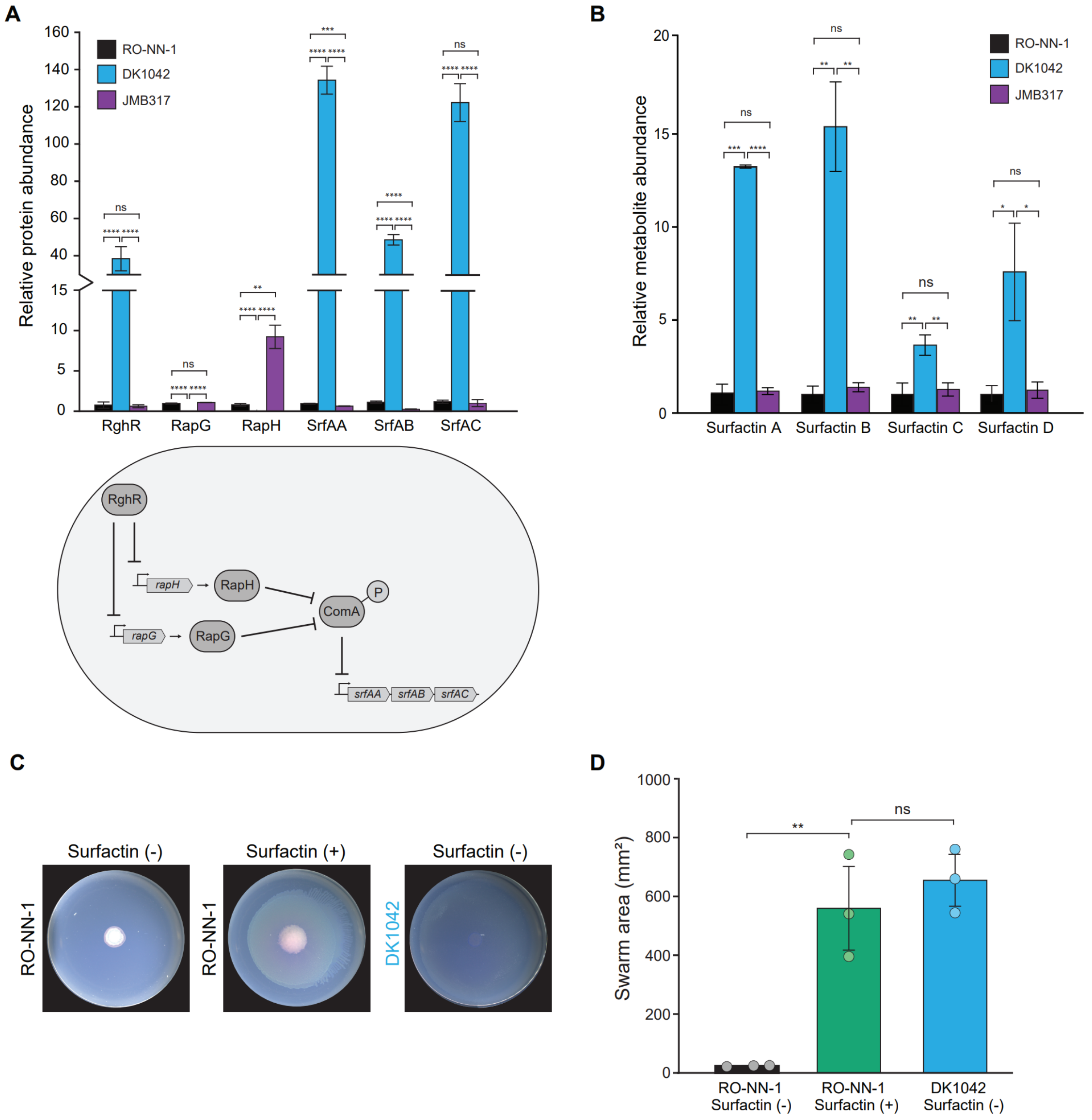

**Fig. S21. Genetic variation gives rise to differences in surfactin production affecting swarming motility in *B. subtilis* RO-NN-1 and DK1042.**

(**A**) Expression of surfactin-producing proteins (SrfAA, SrfAB, and SrfAC), RghR, RapG and RapH was monitored through global proteomics of *B. subtilis* RO-NN-1, DK1042 and JMB317 (DK1042 derivative strain containing the two candidate swarming motility causal genetic variants (Fig. 3E)). (**B**) Surfactin production was measured in the cell pellet from DK1042, RO-NN-1 and JMB317 swarms after 5h of incubation. (**C**) Supplementation of surfactin to the agar rescues the swarming-deficient phenotype of strain RO-NN-1. Comparison of swarming motility of *B. subtilis* DK1042 and RO-NN-1 with and without surfactin supplementation over the course of 5h. Representative data from three biological experiments is shown. (**D**) Areas covered by the DK1042 and RO-NN-1 swarms with and without surfactin supplementation measured after 5h of incubation. The data were analyzed by means of two-tailed Student's *t* test and represents the average results of three biological replicates. Error bars indicate standard deviation. **: $p$ value < 0.01, ns: not significant.

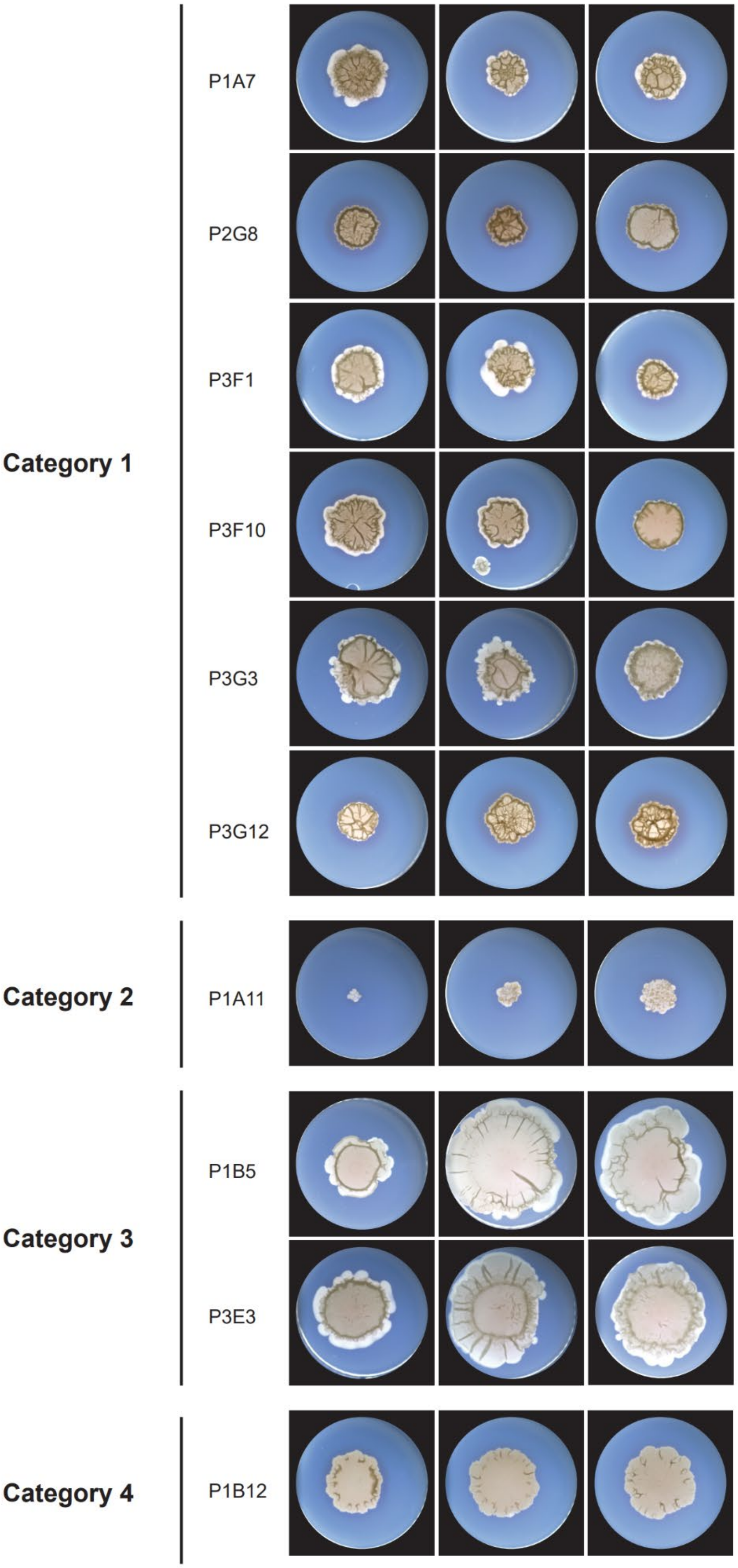
Category 1
P1A7
P2G8
P3F1
P3F10
P3G3
P3G12
Category 2
P1A11
Category 3
P1B5
P3E3
Category 4
P1B12

Continued

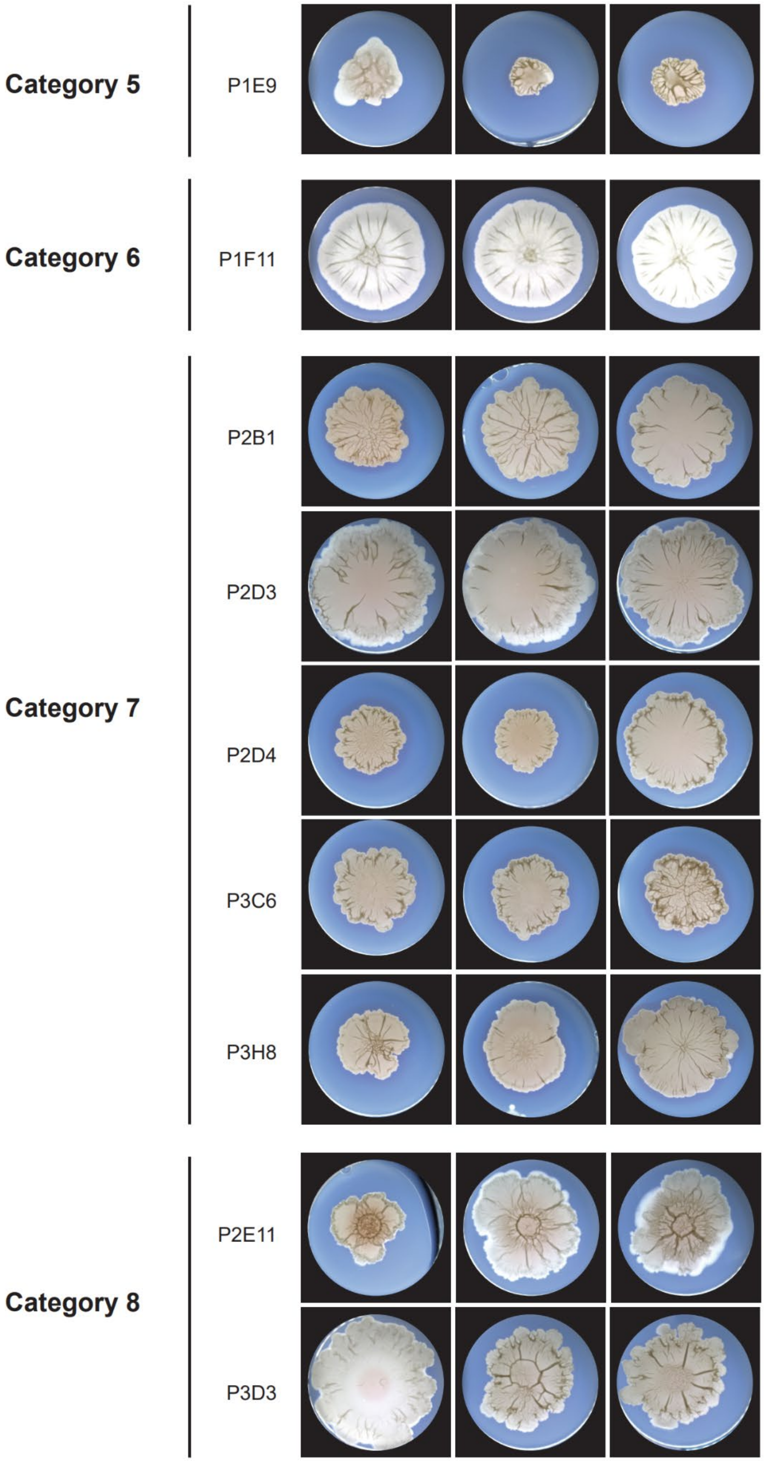
Category 5
P1E9
Category 6
P1F11
Category 7
P2B1
P2D3
P2D4
P3C6
P3H8
Category 8
P2E11
P3D3

Continued

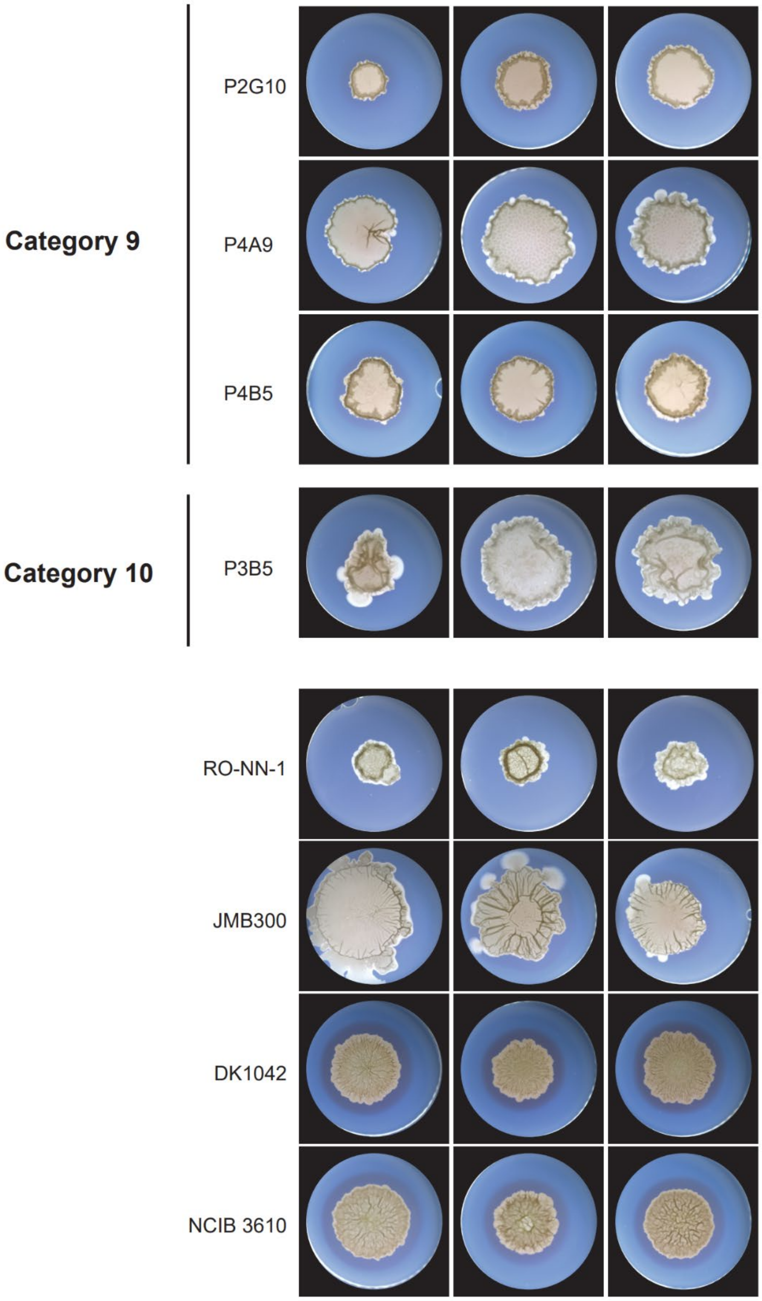

**Fig. S22. Colony morphology varies across a panel of the most genetically diverse *B. subtilis* shuffled strains from the QTL population.**

The 23 most genetically diverse shuffled strains, *B. subtilis* RO-NN-1, JM300, DK1042 and NCIB 3610 were grown on biofilm-promoting minimal agar for 72h. Colonies from three independent biological experiments are shown. The colonies of the shuffled strains were categorized based on their morphology into 10 groups.

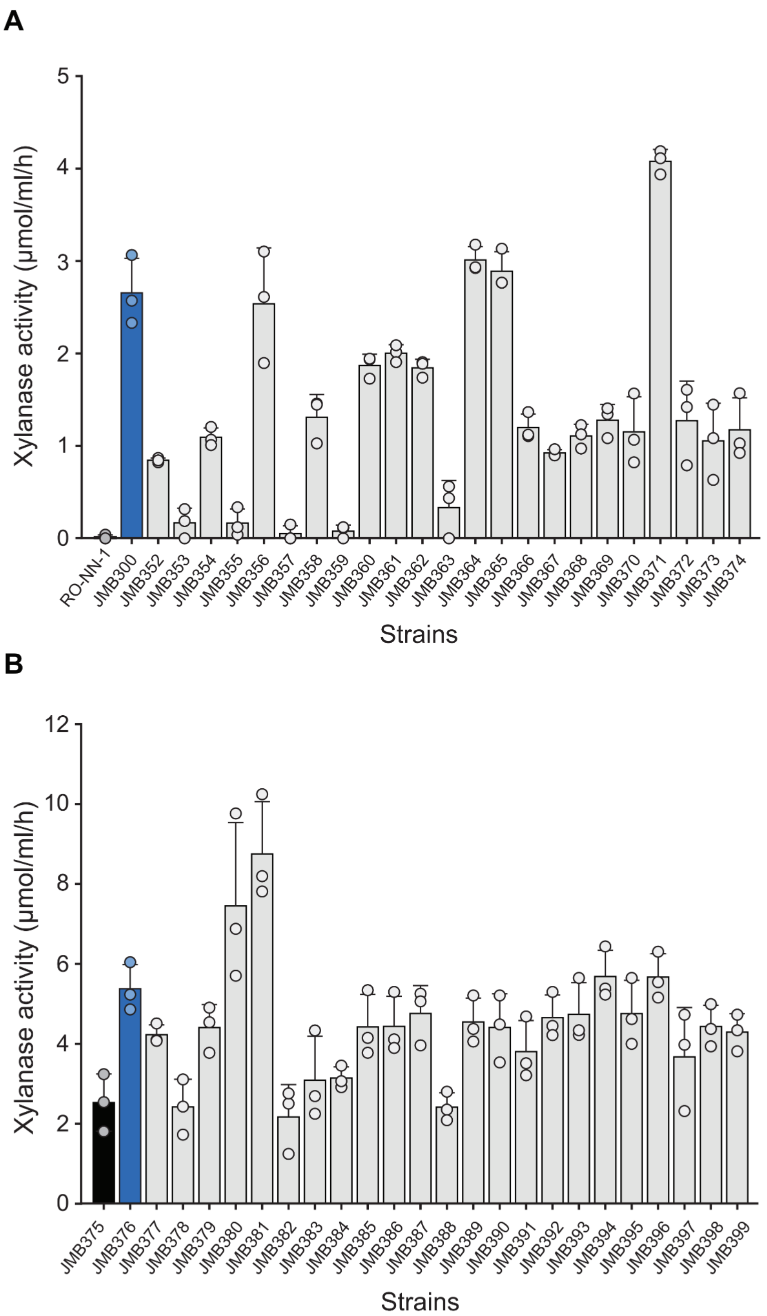

**Fig. S23. Xylanase activity varies across a panel of the most genetically diverse shuffled *B. subtilis* strains from the QTL population.**

(**A**) Intrinsic xylanase activity was measured across the 23 most diverse shuffled strains (JMB352-374), *B. subtilis* RO-NN-1 and JMB300. (**B**) Xylanase activity was measured across the 23 most diverse shuffled strains (JMB352-374), *B. subtilis* RO-NN-1 and JMB300 harboring pJM607 without induction. The data represents the mean results of three biological replicates. Error bars represent standard deviation.

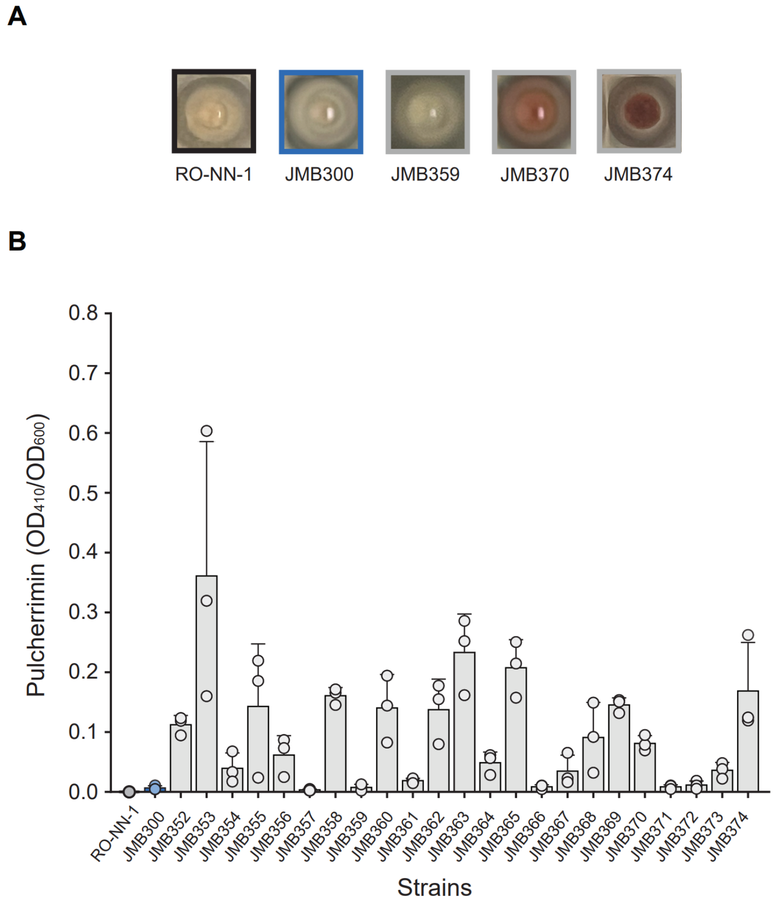

**Fig. S24. Pulcherrimin production varies across a panel of the most genetically diverse *B. subtilis* shuffled strains from the QTL population.**
(**A**) Representative images of cell pellets obtained after harvesting cultures from *B. subtilis* RO-NN-1, JMB300 and three shuffled strains (JMB359, JMB370 and JMB374) grown for 48h in modified LBGM medium supplemented with 0.2 mM $FeCl_3$. Pulcherrimin co-precipitates with the cells after centrifugation. (**B**) Quantification of pulcherrimin production in the 23 most genetically diverse shuffled strains, *B. subtilis* RO-NN-1 and JMB300. Data represents the mean results of three biological replicates with two technical replicates each. Error bars indicate standard deviation.

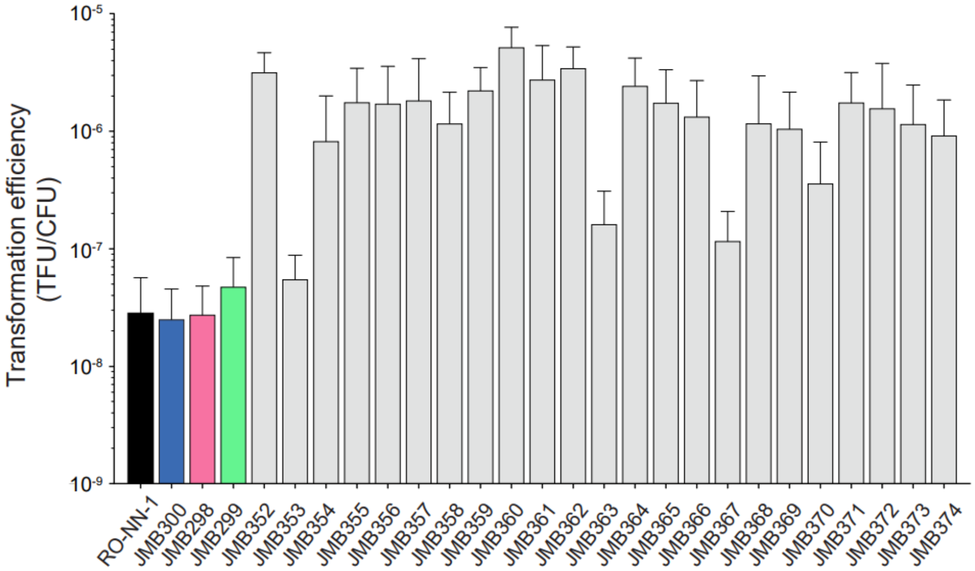

**Fig. S25. Transformation efficiency varies across a panel of the most genetically diverse *B. subtilis* shuffled strains from the QTL population.**

Cells of the 23 most genetically diverse shuffled strains, *B. subtilis* RO-NN-1, JMB298, JMB299, and JM300 were transformed with the replicative plasmid pIDV40 via natural competency. Transformation efficiency was calculated by dividing the number of transformants (TFU) by the total number of viable cells (CFU) per milliliter.

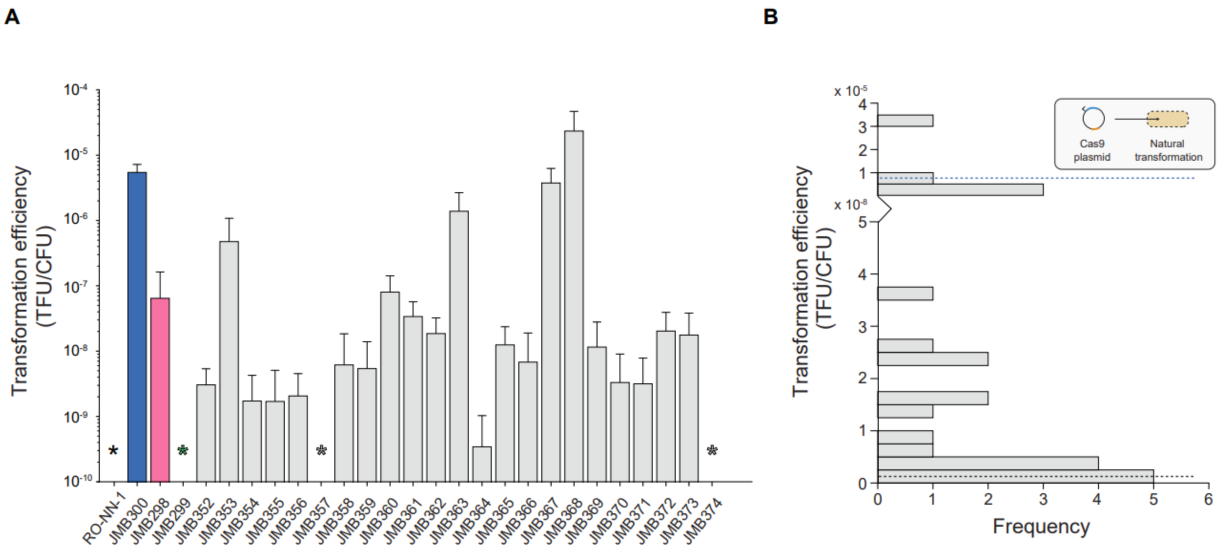

**Fig. S26. Transformation efficiency with a Cas9 plasmid varies across a panel of the most genetically diverse *B. subtilis* shuffled strains from the QTL population.**
(**A**) Cells of the 23 most genetically diverse shuffled strains, *B. subtilis* RO-NN-1, JMB298, JMB299, and JM300 were transformed with pJOE9658.1 (*31*) via natural competency. Transformation efficiency was calculated by dividing the number of transformants (TFU) by the total number of viable cells (CFU) per milliliter. Strains in which no transformants were detected are indicated by asterisks. (**B**) A histogram of the transformation efficiencies is shown. Black and dark blue dashed lines indicate phenotypic data from RO-NN-1 and JM300, respectively.

**Table S1. Strains used in this study.**

| Strain | Genotype | Reference | Figures |
|---|---|---|---|
| *Bacillus subtilis* subsp. *subtilis* RO-NN-1 | Wild type | (*1*, *2*) | 1-4, |
| *Bacillus subtilis* subsp. *subtilis* NCIB3610 | Wild type | (*3*, *4*) | S22 |
| *Bacillus subtilis* DK1042 | NCIB3610 *comI*$^{Q12L}$ | (*5*) | 1-3, |
| *B. subtilis* DSM 1970 | Wild type | | S16 |
| *B. spizizenii* TU-B-10 | Wild type | (*1*, *7*, *8*) | S5, S16 |
| *B. altitudinis* S3 | Wild type | | S16 |
| *B. velezensis* S10-2-W | Wild type | (*46*) | S16 |
| *B. velezensis* S10-2-Y | Wild type | | S16 |
| *B. velezensis* FZB42 | Wild type | (*9*, *10*) | S16 |
| *B. velezensis* GB03 | Wild type | (*11*) | S16 |
| *B. velezensis* S4 | Wild type | (*46*) | S16 |
| *B. velezensis* S141 | Wild type | (*47*) | S16 |
| *Bacillus sp*. BK100 | Wild type | (*48*) | S16 |
| BKK34900 | 168 Δ*hisB*::*kan* | (*6*) | |
| BKE13180 | 168 Δ*metE*::*erm* | (*6*) | |
| JMB1 | RO-NN-1 Δ*hisB*::*kan* | (*12*) | S1 |
| JMB3 | RO-NN-1 Δ*metE*::*erm* | (*12*) | S1 |
| JMB60 | RO-NN-1 Δ*hisB*::*kan* Δ*metE*::*erm* | (*12*) | |
| JMB90 | DK1042 Δ*metE*::*erm* | This study | |
| JMB91 | DK1042 Δ*hisB*::*kan* Δ*metE*::*erm* | This study | |
| JMB92 | DK1042 Δ*hisB*::*erm* | This study | |
| JMB298 | RO-NN-1 Δ*hsdR* | (*12*) | |
| JMB299 | RO-NN-1 Δ*mrr* | (*12*) | |
| JMB241-JMB264 | S1 Δ*metE*::*erm* pool x DK1042 Δ*hisB*::*kan* ➘ S2 *hisB*$^+$ *metE*$^+$ | This study | 1G, S1 |
| JMB300 | NCIB 3610 lacking pBS32 | This study | 4 |

| | | |
|---|---|---|
| JMB301 | RO-NN-1 Δ*gerA*::*kan* | This study |
| JMB302 | RO-NN-1 nt 3,188,758- nt 3,189,735 replaces<br>DK1042 nt 3,391,225- nt 3,392,202<br>(*gerAA* nt 429- *gerAA* nt 1406) | This study |
| JMB303 | RO-NN-1 nt 3,188,407- nt 3,190,148 replaces<br>DK1042 nt 3,390,874- nt 3,392,615<br>(*gerAA* nt 78- *gerAB* nt 402) | This study |
| JMB304 | RO-NN-1 nt 3,188,758- nt 3,190,580 replaces<br>DK1042 nt 3,391,225- nt 3,393,047<br>(*gerAA* nt 429- *gerAB* nt 834) | This study |
| JMB305 | RO-NN-1 nt 3,189,323- nt 3,190,458 replaces<br>DK1042 nt 3,391,790- nt 3,392,925<br>(*gerAA* nt 994- *gerAB* nt 712) | This study |
| JMB306 | RO-NN-1 nt 3,188,869- nt 3,190,070 replaces<br>DK1042 nt 3,391,336- nt 3,392,537<br>(*gerAA* nt 540- *gerAB* nt 324) | This study |
| JMB307 | RO-NN-1 nt 3,188,471- nt 3,189,461 replaces<br>DK1042 nt 3,390,938- nt 3,391,928<br>(*gerAA* nt 142- *gerAA* nt 1,132) | This study |
| JMB308 | RO-NN-1 nt 3,188,407- nt 3,189,580 replaces<br>DK1042 nt 3,390,874- nt 3,392,047<br>(*gerAA* nt 78- *gerAA* nt 1,251)<br><br>RO-NN-1 nt 3,189,661- nt 3,189,735 replaces<br>DK1042 nt 3,392,128- nt 3,392,202<br>(*gerAA* nt 1,332- *gerAA* nt 1,406) | This study |
| JMB309 | RO-NN-1 nt 3,188,758- nt 3,189,460 replaces<br>DK1042 nt 3,391,225- nt 3,391,928<br>(*gerAA* nt 429- *gerAA* nt 1,132) | This study |
| JMB310 | RO-NN-1 nt 3,190,064- nt 3,190,580 replaces<br>DK1042 nt 3,392,531- nt 3,393,047<br>(*gerAB* nt 318- *gerAB* nt 834) | This study |
| JMB311 | RO-NN-1 nt 3,190,070- nt 3,190,702 replaces<br>DK1042 nt 3,392,537- nt 3,393,169 | This study |

| | | |
|---|---|---|
| | (*gerAB* nt 324- *gerAB* nt 956) | |
| JMB312 | RO-NN-1 nt 3,190,230- nt 3,190,727 replaces DK1042 nt 3,392,697- nt 3,393,194 (*gerAB* nt 484- *gerAB* nt 981) | This study |
| JMB313 | RO-NN-1 nt 3,190,230- nt 3,190,580 replaces DK1042 nt 3,392,697- nt 3,393,047 (*gerAB* nt 484- *gerAB* nt 834) | This study |
| JMB314 | RO-NN-1 nt 3,190,064- nt 3,190,607 replaces DK1042 nt 3,392,531- nt 3,393,074 (*gerAB* nt 318- *gerAB* nt 861) | This study |
| JMB315 | RO-NN-1 nt 3,190,230- nt 3,190,550 replaces DK1042 nt 3,392,697- nt 3,393,017 (*gerAB* nt 484- *gerAB* nt 804) | This study |
| JMB316 | RO-NN-1 nt 3,189,223- nt 3,191,809 replaces DK1042 nt 3,391,690- nt 3,394,276 (*gerAA* nt 894- *gerAC* nt 969) | This study |
| JMB317 | RO-NN-1 nt 3,189,572- nt 3,190,150 replaces DK1042 nt 3,392,039- nt 3,392,615 (*gerAA* nt 1,243- *gerAB* nt 404)<br><br>RO-NN-1 nt 3,190,290- nt 3,190,355 replaces DK1042 nt 3,392,757- nt 3,392,822 (*gerAB* nt 544- *gerAB* nt 609)<br><br>RO-NN-1 nt 3,190,403- nt 3,190,436 replaces DK1042 nt 3,392,870- nt 3,392,903 (*gerAB* nt 657- *gerAB* nt 690)<br><br>RO-NN-1 nt 3,190,511- nt 3,191,224 replaces DK1042 nt 3,392,978- nt 3,393,691 (*gerAB* nt 765- *gerAC* nt 384) | This study |
| JMB318 | RO-NN-1 nt 3,189,572- nt 3,191,924 replaces DK1042 nt 3,392,039- nt 3,394,391 (*gerAA* nt 1,243- *gerAC* nt 1,084) | This study |
| JMB319 | RO-NN-1 nt 3,189,572- nt 3,190,458 replaces DK1042 nt 3,392,039- nt 3,392,925 | This study |

| | | |
|---|---|---|
| | (*gerAA* nt 1,243- *gerAB* nt 712) | |
| JMB320 | RO-NN-1 nt 3,190,867- nt 3,191,924 replaces<br>DK1042 nt 3,393,334- nt 3,394,391<br>(*gerAC* nt 27- *gerAC* nt 1,084) | This study |
| JMB321 | RO-NN-1 nt 3,188,471- nt 3,191,315 replaces<br>DK1042 nt 3,390,938- nt 3,393,818<br>(*gerAA* nt 42- *gerAC* nt 511) | This study |
| JMB322 | RO-NN-1 nt 3,190,511- nt 3,191,809 replaces<br>DK1042 nt 3,392,987- nt 3,394,276<br>(*gerAB* nt 765- *gerAC* nt 969) | This study |
| JMB323 | RO-NN-1 nt 3,189,661- nt 3,191,086 replaces<br>DK1042 nt 3,392,128- nt 3,393,557<br>(*gerAA* nt 1,332- *gerAC* nt 246)<br><br>RO-NN-1 nt 3,191,146- nt 3,191,695 replaces<br>DK1042 nt 3,393,613- nt 3,394,162<br>(*gerAC* nt 306- *gerAC* nt 855)<br><br>RO-NN-1 nt 3,191,732- nt 3,191,809 replaces<br>DK1042 nt 3,394,200- nt 3,394,276<br>(*gerAC* nt 892- *gerAC* nt 969) | This study |
| JMB324 | RO-NN-1 nt 3,191,001- nt 3,191,434 replaces<br>DK1042 nt 3,393,468- nt 3,393,901<br>(*gerAC* nt 161- *gerAC* nt 594) | This study |
| JMB325 | RO-NN-1 nt 3,190,727- nt 3,191,536 replaces<br>DK1042 nt 3,393,194- nt 3,394,003<br>(*gerAB* nt 981- *gerAC* nt 594) | This study |
| JMB326 | RO-NN-1 nt 3,190,927- nt 3,191,248 replaces<br>DK1042 nt 3,393,394- nt 3,393,715<br>(*gerAC* nt 87- *gerAC* nt 408)<br><br>RO-NN-1 nt 3,191,284- nt 3,191,924 replaces<br>DK1042 nt 3,393,751- nt 3,394,391<br>(*gerAB* nt 444- *gerAC* nt 1084) | This study |
| JMB327 | RO-NN-1 nt 3,190,867- nt 3,191,782 replaces<br>DK1042 nt 3,393,334- nt 3,394,249 | This study |

| | | |
|---|---|---|
| | (*gerAC* nt 27- *gerAC* nt 942) | |
| JMB328 | RO-NN-1 nt 3,191,162- nt 3,191,209 replaces DK1042 nt 3,393,629- nt 3,393,676 (*gerAC* nt 322- *gerAC* nt 369)<br><br>RO-NN-1 nt 3,191,234- nt 3,191,924 replaces DK1042 nt 3,393,701- nt 3,394,391 (*gerAC* nt 394- *gerAC* nt 1084) | This study |
| JMB329 | RO-NN-1 nt 3,191,001- nt 3,191,086 replaces DK1042 nt 3,393,468- nt 3,393,553 (*gerAC* nt 161- *gerAC* nt 246)<br><br>RO-NN-1 nt 3,191,146- nt 3,191,476 replaces DK1042 nt 3,393,613- nt 3,393,943 (*gerAC* nt 306- *gerAC* nt 636) | This study |
| JMB330 | RO-NN-1 nt 3,391,035- nt 3,191,564 replaces DK1042 nt 3,393,502- nt 3,394,031 (*gerAC* nt 195- *gerAC* nt 724) | This study |
| JMB331 | RO-NN-1 nt 3,391,146- nt 3,191,924 replaces DK1042 nt 3,393,613- nt 3,394,391 (*gerAC* nt 306- *gerAC* nt 1,084) | This study |
| JMB332 | RO-NN-1 nt 3,188,407- nt 3,191,104 replaces DK1042 nt 3,390,874- nt 3,393,571 (*gerAA* nt 78- *gerAC* nt 264) | This study |
| JMB333 | RO-NN-1 nt 3,188,407- nt 3,191,924 replaces DK1042 nt 3,390,874- nt 3,394,391 (*gerAA* nt 78- *gerAC* nt 1,084) | This study |
| JMB334 | RO-NN-1 nt 3,253,390- nt 3,254,911 replaces DK1042 nt 3,452,597- nt 3,454,118 (*rnr* nt 99 - *rnr* nt 1,620) | This study |
| JMB335 | RO-NN-1 nt 3,252,876-nt 3,254,704 replaces DK1042 nt 3,452,084- nt 3,454,118 (*rnr* nt 99 - *rnr* nt 2,133) | This study |
| JMB336 | RO-NN-1 nt 3,252,787- nt 3,255,535 replaces DK1042 nt 3,451,994- nt 3,454,742 | This study |

| | | |
|---|---|---|
| | (*estA* nt 234- *rnr* nt 2,223) | |
| JMB337 | RO-NN-1 nt 3,252,877- nt 3,257,184 replaces<br>DK1042 nt 3,452,084- nt 3,456,391<br>(*yvzC* nt 139- *rnr* nt 2,133) | This study |
| JMB338 | RO-NN-1 nt 3,252,877- nt 3,257,184 replaces<br>DK1042 nt 3,452,084- nt 3,456,391<br>(*yvzC* nt 139- *rnr* nt 2,133)<br>RO-NN-1 nt 3,260,600 - nt 3,263,518 replaces<br>DK1042 nt 3,459,816 - nt 3,462,064 (*opuBB* nt 39 - 4 bp downstream of *opuBD*) | This study |
| JMB339 | RO-NN-1 nt 3,252,877- nt 3,257,184 replaces<br>DK1042 nt 3,452,084- nt 3,456,391<br>(*yvzC* nt 139- *rnr* nt 2,133)<br><br>RO-NN-1 nt 3,259,739- nt 3,260,640 replaces<br>DK1042 3,458,955- nt 3,459,856<br>(*yvaQ* nt 876- *opuBD* nt 645)<br><br>RO-NN-1 nt 3,260,723- nt 3,260,772 replaces<br>DK1042 nt 3,459,931- nt 3,459,990 (*opuBD* nt 511- *opuBD* nt 570)<br><br>RO-NN-1 nt 3,261,065- nt 3,263,821 replaces<br>DK1042 nt 3,460,273- nt 3,463,037<br>(*opuBA* nt 228- *opuBD* nt 228) | This study |
| JMB340 | RO-NN-1 nt 3,252,877- nt 3,257,184 replaces<br>DK1042 nt 3,452,084- nt 3,456,391<br>(*yvzC* nt 139- *rnr* nt 2,133)<br><br>RO-NN-1 nt 3,259,268- nt 3,260,540 replaces<br>DK1042 nt 3,458,484- nt 3,459,755<br>(*yvaQ* nt 405- *yvaQ* nt 1,677)<br><br>RO-NN-1 nt 3,260,608- nt 3,263,363 replaces<br>DK1042 nt 3,459,816- nt 3,462,579<br>(36 bp downstream of *yvaQ*- *opuBA* nt 686) | This study |
| JMB341 | RO-NN-1 nt 3,252,877- nt 3,257,184 replaces<br>DK1042 nt 3,452,084- nt 3,456,391<br>(*yvzC* nt 139- *rnr* nt 2,133)<br><br>RO-NN-1 nt 3,257,747- nt 3,259,025 replaces<br>DK1042 nt 3,456,962- nt 3,458,241<br>(*rghRB* nt 285 - *yvaQ* nt 162) | This study |

| | | |
|---|---|---|
| | RO-NN-1 nt 3,259,745- nt 3,263,686 replaces<br>DK1042 nt 3,458,955- nt 3,462,887<br>(*yvaQ* nt 881- *opuBA* nt 378)<br><br>RO-NN-1 nt 3,264,041- nt 3,264,091 replaces<br>DK1042 nt 3,463,257- nt 3,463,307<br>(43 bp upstream of *opuBA* - *opuBA* nt 8) | |
| JMB342 | RO-NN-1 nt 3,252,877- nt 3,257,184 replaces<br>DK1042 nt 3,452,084- nt 3,456,391<br>(*yvzC* nt 139- *rnr* nt 2,133)<br><br>RO-NN-1 nt 3,258,193- nt 3,260,297 replaces<br>DK1042 3,457,405- nt 3,459,501<br>(*rghRB* nt 292- *yvaQ* nt 1434)<br><br>RO-NN-1 nt 3,260,679- nt 3,263,367 replaces<br>DK1042 nt 3,459,892- nt 3,462,578<br>(*opuBA* nt 682- *opuBD* nt 606) | This study |
| JMB343 | RO-NN-1 nt 3,252,292- nt 3,257,747 replaces<br>DK1042 nt 3,451,499- nt 3,456,962<br>(*smpB* nt 234- *rghRA* nt 282) | This study |
| JMB344 | RO-NN-1 nt 3,252,787- nt 3,254,911 replaces<br>DK1042 nt 3,451,994- nt 3,454,118<br>(*rnr* nt 99- *rnr* nt 2,223)<br><br>RO-NN-1 nt 3,255,535- nt 3,256,708 replaces<br>DK1042 3,454,742- nt 3,455,915<br>(*estA* nt 234- *yvaM* nt 430)<br><br>RO-NN-1 nt 3,256,848- nt 3,258,045 replaces<br>DK1042 nt 3,456,055- nt 3,457,261<br>(*yvaM* nt 570- *rghRB* nt 144) | This study |
| JMB345 | RO-NN-1 nt 3,254,113-nt 3,257,480 replaces<br>DK1042 nt 3,453,320- nt 3,456,695<br>(*rnr* nt 897- *rgrRA* nt 15) | This study |
| JMB346 | RO-NN-1 nt 3,225,511- nt 3,257,480 replaces<br>DK1042 3,454,718- nt 3,456,695<br>(*estA* nt 258- *rghRA* nt 15) | This study |

| | | |
|---|---|---|
| | RO-NN-1 nt 3,252,787- nt 3,253,042<br>replaces<br>DK1042 nt 3,451,994- nt 3,452,249<br>(*rnr* nt 1968- *rnr* nt 2,223) | |
| JMB347 | RO-NN-1 nt 3,255,511-nt 3,258,045<br>replaces<br>DK1042 nt 3,454,718- nt 3,457,261<br>(*estA* nt 258- *rghRB* nt 144) | This study |
| JMB348 | RO-NN-1 nt 3,256,683- nt 3,258,201<br>replaces<br>DK1042 nt 3,455,890- nt 3,457,417<br>(*yvaM* nt 405- *rghRB* nt 300) | |
| JMB349 | RO-NN-1 nt 3,256,050- nt 3,257,867<br>replaces<br>DK1042 nt 3,455,257- nt 3,457,082<br>(*secG* nt 81- *rghRA* nt 401) | This study |
| JMB350 | RO-NN-1 nt 3,256,863<br>replaces<br>DK1042 nt 3,256,863<br>(*yvaM* nt 585)<br><br>RO-NN-1 nt 3,257,348- nt 3,257,435<br>replaces<br>DK1042 nt 3,456,555- nt 3,456,650<br>(126 bp upstream of *rghRA*- 31 bp upstream of *rghRA*) | This study |
| JMB351 | RO-NN-1 nt 3,256,740<br>replaces<br>DK1042 nt 3,455,947<br>(*yvaM* nt 462)<br><br>RO-NN-1 nt 3,257,340- nt 3,257,480<br>replaces<br>DK1042 nt 3,456,547- nt 3,456,695<br>(134 bp upstream of *rghRA*- *rghRA* nt 15) | This study |
| JMB352 | *B. subtilis* P1A7 Δ*hisB* | This study |
| JMB353 | *B. subtilis* P1A11 Δ*hisB* | This study |
| JMB354 | *B. subtilis* P1B5 Δ*hisB* | This study |
| JMB355 | *B. subtilis* P1B12 Δ*hisB* | This study |
| JMB356 | *B. subtilis* P1E9 Δ*hisB* | This study |
| JMB357 | *B. subtilis* P1F11 Δ*hisB* | This study |
| JMB358 | *B. subtilis* P2B1 Δ*hisB* | This study |

| | | |
|---|---|---|
| JMB359 | *B. subtilis* P2D3 Δ*hisB* | This study |
| JMB360 | *B. subtilis* P2D4 Δ*hisB* | This study |
| JMB361 | *B. subtilis* P2E11 Δ*hisB* | This study |
| JMB362 | *B. subtilis* P2G8 Δ*hisB* | This study |
| JMB363 | *B. subtilis* P2G10 Δ*hisB* | This study |
| JMB364 | *B. subtilis* P3B5 Δ*hisB* | This study |
| JMB365 | *B. subtilis* P3C6 Δ*hisB* | This study |
| JMB366 | *B. subtilis* P3D3 Δ*hisB* | This study |
| JMB367 | *B. subtilis* P3E3 Δ*hisB* | This study |
| JMB368 | *B. subtilis* P3F1 Δ*hisB* | This study |
| JMB369 | *B. subtilis* P3F10 Δ*metE* | This study |
| JMB370 | *B. subtilis* P3G3 Δ*metE* | This study |
| JMB371 | *B. subtilis* P3G12 Δ*metE* | This study |
| JMB372 | *B. subtilis* P3H8 Δ*metE* | This study |
| JMB373 | *B. subtilis* P4A9 Δ*metE* | This study |
| JMB374 | *B. subtilis* P4B5 Δ*metE* | This study |
| JMB375 | *B. subtilis* RO-NN-1 carrying pJM607 | This study |
| JMB376 | JMB300 carrying pJM607 | This study |
| JMB377 | *B. subtilis* P1A7 Δ*hisB* carrying pJM607 | This study |
| JMB378 | *B. subtilis* P1A11 Δ*hisB* carrying pJM607 | This study |
| JMB379 | *B. subtilis* P1B5 Δ*hisB* carrying pJM607 | This study |
| JMB380 | *B. subtilis* P1E9 Δ*hisB* carrying pJM607 | This study |
| JMB381 | *B. subtilis* P1F11 Δ*hisB* carrying pJM607 | This study |
| JMB382 | *B. subtilis* P2B1 Δ*hisB* carrying pJM607 | This study |
| JMB383 | *B. subtilis* P2D3 Δ*hisB* carrying pJM607 | This study |
| JMB384 | *B. subtilis* P2D4 Δ*hisB* carrying pJM607 | This study |
| JMB385 | *B. subtilis* P2E11 Δ*hisB* carrying pJM607 | This study |
| JMB386 | *B. subtilis* P2G8 Δ*hisB* carrying pJM607 | This study |

| | | |
|---|---|---|
| JMB387 | *B. subtilis* P2G10 Δ*hisB* carrying pJM607 | This study |
| JMB388 | *B. subtilis* P3B5 Δ*hisB* carrying pJM607 | This study |
| JMB389 | *B. subtilis* P3C6 Δ*hisB* carrying pJM607 | This study |
| JMB390 | *B. subtilis* P3D3 Δ*hisB* carrying pJM607 | This study |
| JMB391 | *B. subtilis* P3E3 Δ*hisB* carrying pJM607 | This study |
| JMB392 | *B. subtilis* P3F1 Δ*hisB* carrying pJM607 | This study |
| JMB393 | *B. subtilis* P3F10 Δ*metE* carrying pJM607 | This study |
| JMB394 | *B. subtilis* P3G3 Δ*metE* carrying pJM607 | This study |
| JMB395 | *B. subtilis* P3G12 Δ*metE* carrying pJM607 | This study |
| JMB396 | *B. subtilis* P3H8 Δ*metE* carrying pJM607 | This study |
| JMB397 | *B. subtilis* P4A9 Δ*metE* carrying pJM607 | This study |
| JMB398 | *B. subtilis* P4B5 Δ*metE* carrying pJM607 | This study |

<insert Table S1 here followed by a page break >

**Table S2. Plasmids used in this study.**

| Plasmid | Genotype | Reference |
| --- | --- | --- |
| pDR244 | $ori_{pACYC}$, spcR, rep $p15A^{ts}$, $P_{PA}$-cre, | (*6*) |
| pJOE9658.1 | $ori_{pUC18}$, kanR, rep $pE194^{ts}$, gRNA, $P_{tetLM}$-tetLM-cas9 | (*31*) |
| pHT254 | P*grac*100-MCS-His, cmR, ampR | (*49*) |
| pJM598 | pJOE9658-sgRNA(pBS32-*repN*) | This study |
| pJM599 | pJOE9658-sgRNA(3610-*gerAA*) | This study |
| pJM600 | pJOE9658-sgRNA(3610-*gerAB*) | This study |
| pJM601 | pJOE9658-sgRNA(3610-*gerAC*) | This study |
| pJM602 | pJOE9658-sgRNA(3610-*gerAC*)-*gerAC*(RO-NN-1) | This study |
| pJM603 | pJOE9658-sgRNA(3610-*rnr*) | This study |
| pJM604 | pJOE9658-sgRNA(3610-*yvaQ*) | This study |
| pJM605 | pJOE9658-sgRNA(3610-*opuBC*) | This study |
| pJM606 | pJOE9658-sgRNA(3610-*rghR*) | This study |
| pJM607 | pHT254-$YmwC^{SP}$-*xylA* | This study |
| pIDV40 | | Del Valle I. *et al.* (in preparation) |

<insert Table S2 here followed by a page break >

**Table S3. Oligonucleotides used in this study.**

| Oligo | Sequence | Used for |
|---|---|---|
| **hisB-FL** | CAATTGCCGGATATAATGTAAAAGCAC | Construction of DK1042 histidine auxotrophic strain |
| **hisB-RL** | ATATGATTGCCGGACCGAGTGAAATC | |
| **metE-FL** | CATGCCTGATCCTTTTAATATTCTTTCTTATTG | Construction of DK1042 methionine auxotrophic strain |
| **metE-RL** | GCTATGAAGAAGAATCATTTCAAAGAAAG | |
| | TATTAGCTAGAGCTGTGC | Deletion of *gerA* in strain RO-NN-1 |
| | tgtatgctatacgaacggtaCAATGAGGTCACCTCTTATC | |
| | gataagaggtgacctcattgTACCGTTCGTATAGCATAC | |
| | cgtgaattaggcggctgctaTCTACCGTTCGTATAATGTATG | |
| | tacattatacgaacggtagaTAGCAGCCGCCTAATTCAC | |
| | GTCATCGGCGAAGCATCG | |
| | TACGATTGGAAAGACGAACTCGGA | pBS32(sgRNA-*repN*) |
| | AAACTCCGAGTTCGTCTTTCCAAT | |
| | TACGAGTCACCATCGAGCTGCTAA | 3610(sgRNA-*gerAA*) |
| | AAACTTAGCAGCTCGATGGTGACT | |
| | TACGTAAAAAAGAGATGGAAACGA | 3610(sgRNA-*gerAB*) |
| | AAACTCGTTTCCATCTCTTTTTTA | |
| | TACGTAAGGTGACGATCAGAACGA | 3610(sgRNA-*gerAC*) |
| | AAACTCGTTCTGATCGTCACCTTA | |

| | | |
|---|---|---|
| | AAGGCCAACGAGGCCTTGGAACAAACAGAGTTTAAGGA | RO-NN-1 *gerAA* repair template and RO-NN-1 *gerAA-gerAC* repair template |
| | AAGGCCTTATTGGCCTTAAGTTTCAGTGGAGTCTGTT | |
| | AAGGCCAACGAGGCCATGAGCCAAAAACAGACTCCAC | RO-NN-1 *gerAB* repair template |
| | AAGGCCTTATTGGCCTCATTTTGCTGTAATCCTCCTTTTG | |
| | AAGGCCAACGAGGCCATGAAAATCCGGATTTTATGTATGT | RO-NN-1 *gerAC* repair template and RO-NN-1 *gerAA-gerAC* repair template |
| | AAGGCCTTATTGGCCCTATTTGTTTGCGCCTTTCGT | |
| | TACGTGTAGCAAACGAAACAGTTG | 3610(sgRNA-*rnr*) |
| | AAACCAACTGTTTCGTTTGCTACA | |
| | TACGAGCCATTTCGTTCATCGTCT | 3610(sgRNA-*yvaQ*) |
| | AAACAGACGATGAACGAAATGGCT | |
| | TACGGCAGCAGGCTTTAATGAACG | 3610(sgRNA-*opuBC*) |
| | AAACCGTTCATTAAAGCCTGCTGC | |
| | TACGCTAAAATATGAGTAACAGTG | 3610(sgRNA-*rghR*) |
| | AAACCACTGTTACTCATATTTTAG | |
